# ATGL-mediated lipolysis is essential for myocellular mitochondrial function and augments PPARδ-induced improvements in mitochondrial respiration

**DOI:** 10.1101/2024.10.31.621255

**Authors:** Anne Gemmink, Tineke van de Weijer, Gert Schaart, Gernot F. Grabner, Esther Kornips, Kèvin Knoops, Rudolf Zechner, Martina Schweiger, Matthijs K.C. Hesselink

## Abstract

Defects in ATGL-mediated myocellular LD lipolysis results in mitochondrial dysfunction of unknown origin, which can be rescued by PPAR agonists. Here we examine whether ATGL-mediated lipolysis is required to maintain mitochondrial network connectivity and function. Moreover, we explored if the functional implications of ATGL deficiency for mitochondrial network dynamics and function can be alleviated by promoting PPARα and/or PPARδ transcriptional activity. To this end, we cultured human primary myotubes from patients with neutral lipid storage disease with myopathy (NLSDM), a rare metabolic disorder caused by a mutation in the PNPLA2 gene. These myotubes possess dysfunctional ATGL and compromised LD lipolysis. In addition, mitochondria-LD contacts, mitochondrial network dynamics, and TMRM intensity were abrogated. Using a humanized ATGL inhibitor in myotubes cultured form healthy donors, revealed similar results. Upon stimulating PPARδ transcriptional activity, mitochondrial respiration improved by more than 50% in human primary myotubes from healthy lean individuals. This increase in respiration was dampened in myotubes with dysfunctional ATGL. Stimulation of PPARδ transcriptional activity had no effect on mitochondria-LD contacts, mitochondrial network dynamics, and TMRM intensity. Our results demonstrate that dysfunctional ATGL results in compromised mitochondrial-LD contacts and mitochondrial dynamics, and that functional ATGL is required to improve mitochondrial respiratory capacity upon stimulation of PPARδ transcriptional activity.

## INTRODUCTION

Neutral lipid storage disease with myopathy (NLSDM) is a rare metabolic myopathy originating from mutations in the PNPLA2 gene that encodes for adipose triglyceride lipase (ATGL) (1–4), a key enzyme for cytosolic lipid droplet (LD) lipolysis (5). In NLSDM patients the defect in triglyceride lipolysis results in abundant ectopic fat storage in multiple organs, including skeletal muscle (1, 2, 4, 6, 7). In addition, NLSDM patients present with, muscle weakness (1, 3, 6, 7), reduced mitochondrial (fat) oxidative capacity (and compromised bioenergetics) (1), insulin resistance (1), and exercise intolerance (2, 3). A link between excess myocellular fat storage and compromised insulin sensitivity is commonly reported (8–10), particularly if the increase in myocellular fat occurs in parallel by a low fat oxidative capacity content and originates from compromised lipolysis rather than increased fat storage capacity (11). How defects in myocellular lipolysis may result in compromised mitochondrial function and its associated clinical symptoms like insulin resistance and myopathy, is however unclear. Given the central stage of compromised mitochondrial fat oxidative capacity in multiple (ageing-related) metabolic disorders like e.g. type 2 diabetes, NLSDM-derived data may help to provide mechanistic insight into the interaction of functional LD lipolysis and mitochondrial function.

Mitochondrial biogenesis and mitochondrial function are under control of peroxisome proliferator-activated receptor (PPAR)-mediated oxidative gene expression, which in turn is known to rely on endogenous fatty-acids as ligands (12). In ATGL deficient mice cardiac mitochondrial function is compromised (13). This is in line with observations in humans with dysfunctional ATGL where a reduced myocellular mitochondrial function is observed (1). In humans with dysfunctional ATGL, the mitochondrial defect can be partly restored upon administration of the pan-PPAR agonist bezafibrate. Treatment with a PPARδ agonist induces oxidative gene expression and mitochondrial biogenesis via induction of PGC1-α (14) along with increased skeletal muscle oxidative capacity (14–16). Interestingly, promoting myocellular lipolytic capacity by overexpressing ATGL, augmented PPARδ activity along with improved mitochondrial biogenesis and oxidative capacity in C2C12 cells (17).

Over the past few years, it has become progressively clear that not only the number and quality of mitochondria in muscle is essential for optimal fat oxidative capacity, but that mitochondrial connectivity (18, 19) and mitochondrial tethering to LDs (18) are determinants of muscle fat oxidative capacity as well. Electron microscopy images of muscle biopsies obtained from patients with dysfunctional ATGL reveals poor mitochondrial network morphology, aberrations in subcellular distribution and disruptions in mitochondrial LD contact sites (6, 7). This suggests a role for ATGL in maintaining mitochondrial network connectivity and fat oxidative capacity.

Using myotubes derived from muscle biopsies of two NLSDM patients, we studied the hypothesis that ATGL deficiency impairs skeletal muscle mitochondrial respiration, network connectivity and mitochondria-LD contacts and will examine if promoting PPARα and/or PPARδ transcriptional activity can abolish the functional implications of ATGL deficiency for mitochondrial respiration and network dynamics. To examine the direct and acute effects of ATGL-mediated lipolysis on mitochondrial connectivity and function, we will use a humanized inhibitor of ATGL (20) in myotubes cultured from healthy donors. In both models, we will examine if exogenous ligands to stimulate oxidative gene expression (PPAR agonists) promote mitochondrial network connectivity and function.

## RESULTS

### Intramyocellular lipid content and mitochondrial abnormalities in biopsies of muscle with dysfunctional ATGL

We previously reported that NLSDM patients exhibit reduced mitochondrial function and *ex vivo* lipid oxidative capacity (1). On electron microscopy images a high intramyocellular lipid content and mitochondrial abnormalities are observed (Suppl. Fig 1). While in healthy skeletal muscle, mitochondria are located at both sides of a z-line at the cross point of two parallel sarcomeres, this is not the case in skeletal muscle with dysfunctional ATGL. Moreover, mitochondria are more often dispersed between the contractile elements and accumulate in certain subcellular fractions when ATGL is deficient (Suppl. Fig 1).

### Dysfunctional ATGL increases intramyocellular lipid content and reduces lipolysis in human primary myotubes

To investigate the mechanisms involved in impaired mitochondrial function upon ATGL deficiency, we isolated and cultured primary myotubes from two NLSDM patients. One patient had a single heterozygous (S-Hz) mutation, while the other patient had a compound heterozygous (C-Hz) mutation (1). First, we examined whether impairments in intracellular lipid degradation were retained in human primary myotubes cultured from these NLSDM patients. ATGL protein levels were lower in myotubes of NLSDM patients (4.95 AU in S-Hz and 3.94 AU in C-Hz), compared to myotubes of healthy lean donors (7.13 (4.87-9.39) AU) (Fig 1A). Besides ATGL, also protein levels of HSL were drastically reduced in myotubes of both NLSDM patients (0.09 and 0.11 for S-Hz and C-Hz, respectively) compared to myotubes of healthy donors (0.30 (0.24-0.37) AU, (Fig 1B)). These differences in lipases mimic the differences observed in the NLSDM patients. Protein abundance of CGI-58 was not different amongst donors (0.23 (0.08-0.38) vs. 0.22 vs. 0.22 for resp. healthy lean, S-Hz and C-Hz, Fig 1C). FA loading increased intracellular lipid content more profoundly in myotubes from the NLSDM patient with the C-Hz mutation (103.6 AU) than in the NLSDM patient with the S-Hz mutation (59.3 AU) or in the healthy lean control (60.0 (29.3-90.7) AU) (Fig 1D-E). In line with these findings, basal lipolysis rates were lower in C-Hz patients (0.03 AU/3h) compared to S-Hz patients (0.18 AU/3h) and healthy lean controls (0.26 (0.12-0.40) AU/3h) (Fig 1F). To confirm that impaired lipid degradation in primary myotubes of NLSDM patients is due to dysfunctional/inactive ATGL we incubated myotubes of healthy lean donors with an ATGL specific inhibitor (NG-497) and examined intramyocellular lipid content. Inhibition of ATGL during an overnight FA-loading resulted in a much higher intramyocellular lipid content (Fig 1G). These results indicate that, similar as skeletal muscle tissue, also cultured primary myotubes of NLSDM patients have reduced lipolysis and accumulate neutral lipids. The acute use of a selective ATGL inhibitor further stresses the notion that these observed differences directly result from dysfunctional/inactive ATGL.

**Figure 1.**
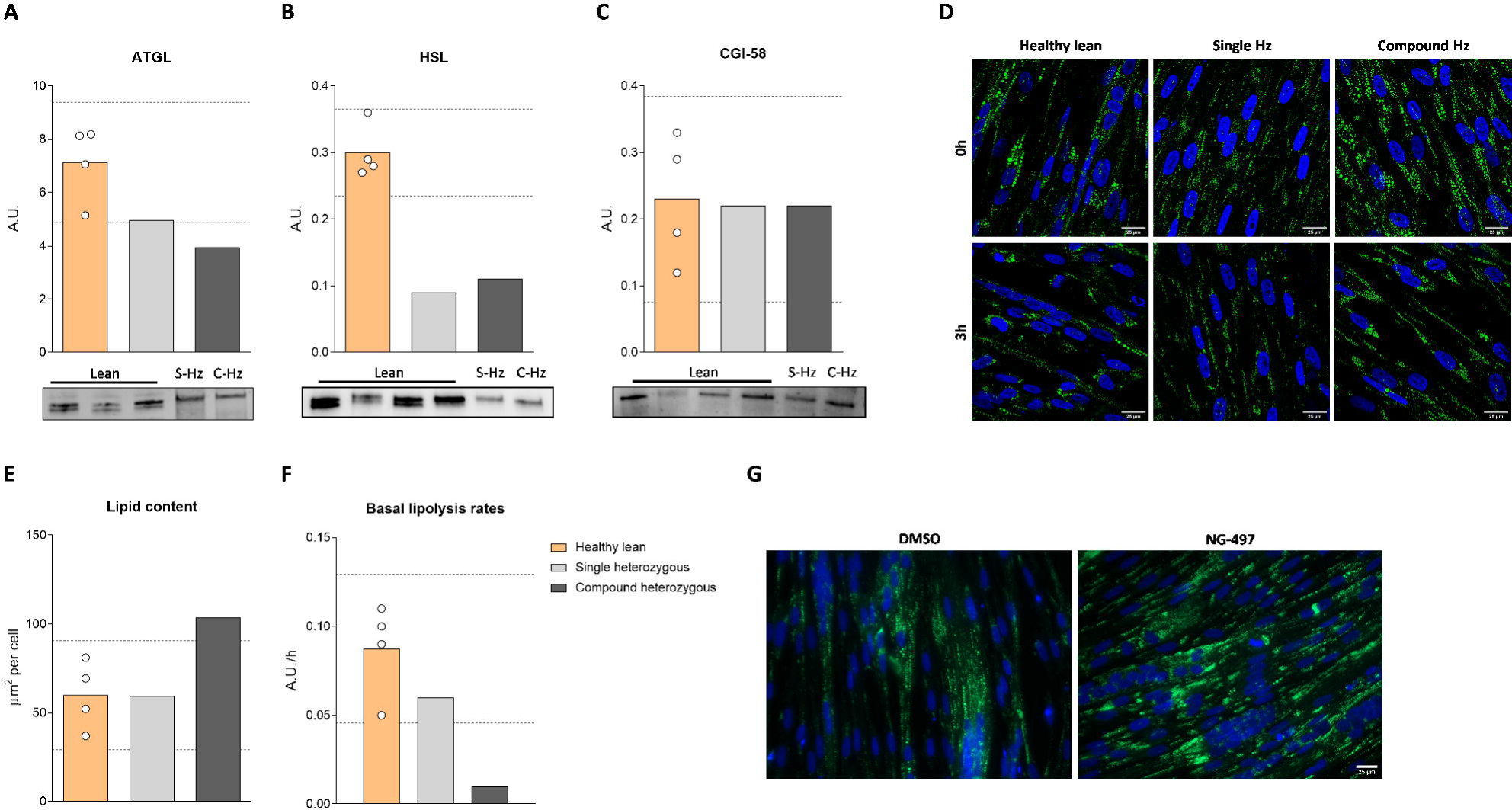
Dysfunctional ATGL increases intramyocellular lipid content and reduces lipolysis in human primary myotubes. (A) ATGL, (B) HSL and (C) CGI-58 protein content. (D) Intramyocellular lipid content after an overnight incubation with 200 μM oleate in myotubes of healthy lean donors and NLSDM patients. LDs and nuclei are in respectively green and blue. (E) Quantification of intramyocellular lipid content. (F) Basal lipolysis rates determined by confocal microscopy. (G) Intramyocellular lipid content upon inhibition of ATGL with NG-497 in myotubes of healthy lean donors. Data are presented as mean, the 95% confidence interval of the healthy lean donors is depicted with the dashed line.

### ATGL deficiency compromises myotube differentiation

NLSDM patients display amongst others with myopathy. Thus, it is of importance to explore if myopathy related defects are also present in primary myotubes of NLSDM patients and if so, if this is due to a reduced ATGL activity. Hence, we monitored the differentiation of myoblasts into myotubes. After differentiation, myotubes of NLSDM patients were smaller compared to myotubes of healthy lean donors, with fewer α-actinin positive cells and lower MF20 abundance (Suppl. Fig 2A). Also, acute ATGL inhibition during differentiation resulted in fewer α-actinin positive myotubes and lower MF20 abundance (Suppl. Fig 2B). In contrast to myotubes of NLSDM patients, ATGL-inhibited myotubes showed a multinuclear phenotype, indicating that ATGL inhibited cells do fuse, but show compromised myotube maturation. Similar to the myotubes of NLSDM patients, also ATGL inhibition by NG-497 during the differentiation process resulted in smaller myotubes, compared to control myotubes. Jointly, these results indicate that compromised ATGL activity impairs proper myotube differentiation.

### Dysfunctional ATGL reduces mitochondrial function

Next, we examined whether dysfunctional ATGL is linked to mitochondrial dysfunction in human primary myotubes. Respiration rates were lower in myotubes derived from NLSDM patients (basal: 1.66 and 2.19, ATP linked: 1.30 and 1.63, and maximal: 5.09 and 6.75 pmol/min/μg protein for S-Hz and C-Hz, respectively), compared to myotubes from healthy lean donors (basal: 3.43 (2.78-4.08), ATP-linked (3.00 (2.46-3.54) and maximal (8.87 (7.31-10.43) pmol/min/μg protein) (Fig 2A-C). Similarly, also inhibition of ATGL activity by NG-497 during the differentiation of myotubes obtained from healthy lean donors caused reduced basal respiration (4.03 (3.08-4.98) (median and range) and 2.83 (2.40-3.25) pmol/min/μg protein for DMSO (control) and NG-497 (inhibitor), respectively p<0.05, Fig 2D) and ATP-linked respiration (3.41 (2.84-3.97) and 2.20 (1.44-2.96) pmol/min/μg protein for DMSO and NG-497, respectively p<0.05, Fig 2E). Maximal respiration, however, was not affected by acute ATGL inhibition (6.99 (3.62-10.36) and 6.57 (4.73-8.41) pmol/min/μg protein for DMSO and NG-497, respectively p=0.663, Fig 2F). To examine whether lower respiration rates were due to lower mitochondrial content or due to mitochondrial dysfunction we measured mitochondrial content on protein and DNA level. Based on the protein abundance of OXPHOS complexes I – V, mitochondrial protein content was similar in all donors with slightly higher content of the complexes II, III, and V in myotubes of NLSDM patients compared to control healthy donors (2.30 (1.82-2.77) vs. 2.19 vs. 2.88 AU for resp. healthy lean, S-Hz and C-Hz, Fig 2G). Similarly, mtDNA copy number computed as the ratio of mitochondrial over nuclear DNA (*ND1/LPL*) was higher in myotubes of NLSDM patients (1588.6 and 1409.6 for S-Hz and C-Hz, respectively) compared to myotubes of healthy lean subjects (935.8 (673.5-1198.1), (Fig 2H). These results indicate that the lower respiratory capacity observed in myotubes with dysfunctional ATGL originates from intrinsic mitochondrial dysfunction and not from reduced mitochondrial content.

**Figure 2.**
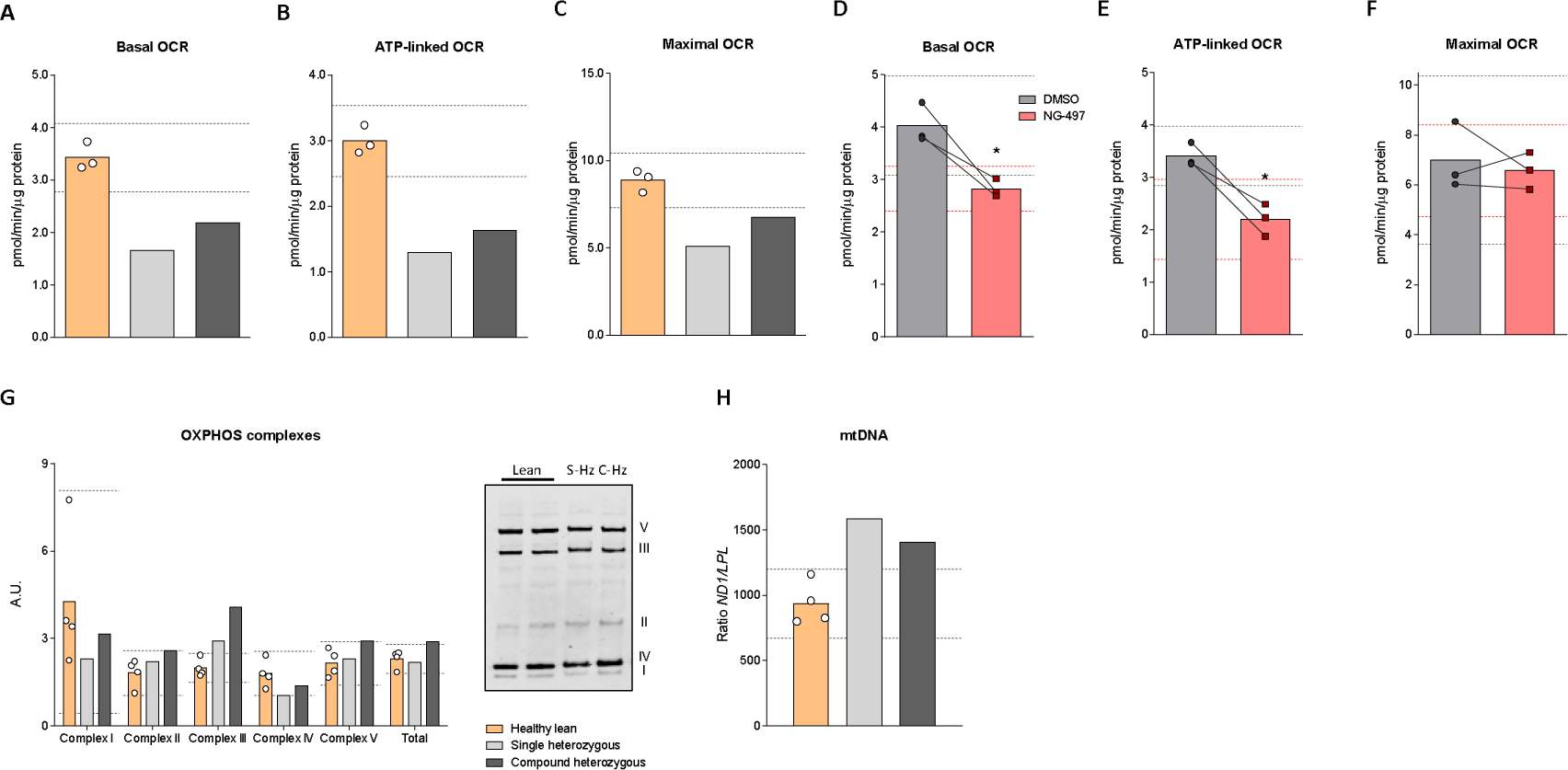
Dysfunctional ATGL reduces mitochondrial function. (A) Basal, (B) ATP-linked and (C) maximal oxygen consumption rates determined with Seahorse in myotubes of healthy lean donors and NLSDM patients. (D) Basal, (E) ATP-linked and (F) maximal oxygen consumption rates in healthy lean donors upon ATGL inhibition with NG-497. (G) Protein content of the different complexes of the mitochondrial respiratory chain. (H) mtDNA content. Data are presented as mean, the 95% confidence interval of the healthy lean donors is depicted with the dashed line.

### ATGL dysfunction alters mitochondrial dynamics

To investigate whether reduced or a lack in ATGL activity affects mitochondrial network integrity we performed live-cell imaging using the mitochondrial membrane potential sensitive dye TMRM. The mitochondrial fragmentation index (MFI) was used as a readout for the level of mitochondrial network fragmentation. We found higher mitochondrial network fragmentation in myotubes of the S-Hz NLSDM patient (1.92) which was exacerbated in myotubes of the C-Hz NLSDM patient (3.00), compared to myotubes of healthy lean donors 1.04 (0.79-1.29) (Fig 3A-B). Intensity of TMRM was lower in myotubes of NLSDM patients (1.64×10^7^ and 1.50×10^7^ AU for S-Hz and C-Hz, respectively) compared to myotubes from healthy lean donors (1.82×10^7^ (1.73×10^7^-1.92×10^7^) AU), (Fig 3A and 3C). Chronic and acute inhibition of ATGL in myotubes from healthy lean donors resulted in severe mitochondrial network fragmentation (Fig 3D, and Suppl. Fig 3B), although this effect was not statistically significant when MFI was used as readout (0.71 (0.52-0.89) vs. 1.09 (0.21-1.97) for DMSO vs. NG-497, p=0.262, Fig 3E). In line with the data obtained from myotubes of NLSDM patients, chronic or acute inhibition of ATGL in myotubes of healthy lean resulted in a significant decrease in intensity of TMRM, compared to DMSO treated controls (1.02×10^7^ (0.79 x10^7^-1.25 x10^7^) and 0.79×10^7^ (0.55 x10^7^-1.02 x10^7^) AU for DMSO and NG-497, respectively, p<0.05, Fig 3F and Suppl. Fig 3B). After a 3.5h wash-out of the ATGL inhibitor, mitochondrial network integrity and TMRM intensity was restored (Suppl. Fig 3A) indicating that ATGL activity acutely and reversibly compromises mitochondrial function.

**Figure 3.**
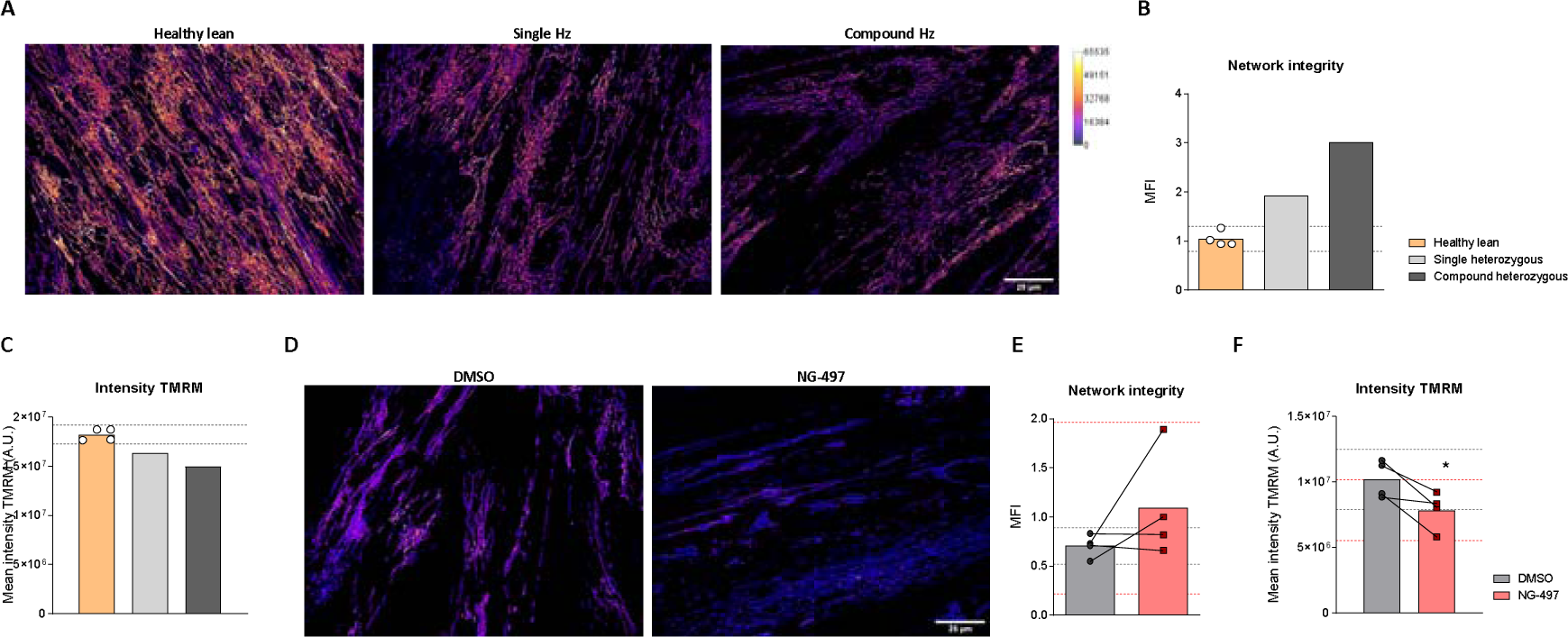
ATGL dysfunction alters mitochondrial dynamics. (A) Representative spinning disk confocal images of myotubes of healthy lean donors and NLSDM patients stained with TMRM depicting network integrity. (B) Quantification of mitochondrial network integrity and (C) intensity TMRM. (D) Spinning disk confocal images of myotubes of healthy lean donors upon ATGL inhibition with NG-497 with TMRM. (E) Quantification of mitochondrial network integrity and (F) intensity TMRM upon inhibition with NG-497. Data are presented as mean, the 95% confidence interval of the healthy lean donors is depicted with the dashed line.

To decipher the mechanisms underlying impaired mitochondrial dynamics in myotubes with compromised ATGL activity we analyzed the expression of different proteins involved in the cycle of mitochondrial network dynamics in myotubes from NLSDM patients. PGC-1α, an important regulator of mitochondrial biogenesis, was lower in myotubes from both NLSDM patients (0.008 and 0.010 AU for S-Hz and C-Hz, respectively) compared to myotubes from healthy lean donors (0.017 (0.009-0.024) AU, Suppl. Fig 4A). Similarly, MFN1, which is involved in mitochondrial fusion was lower in myotubes of both NLSDM patients (0.10 and 0.11 AU, for S-Hz and C-Hz) compared to healthy lean donors (0.20 (0.12-0.29) AU, (Suppl. Fig 4B), while MFN2 and OPA1 were only modestly lower (MFN2: 0.035 (0.028-0.041), 0.027 and 0.033 AU for healthy lean, S-Hz and C-Hz respectively, Fig 4C) or not different (OPA1: 0.11 (0.08-0.13), 0.10 and 0.11 AU or healthy lean, S-Hz and C-Hz respectively, Fig 4D) in myotubes of NLSDM patients. Protein expression of DRP1 and FIS1, important for mitochondrial fission, was also lower in myotubes from both NLSDM patients compared to myotubes from healthy lean controls (DRP1: 0.024 (0.019-0.028), 0.008 and 0.010 AU for healthy lean, S-Hz and C-Hz, respectively; FIS1: 0.090 (0.058-0.122), 0.044, and 0.070 AU for healthy lean, S-Hz and C-Hz, respectively) (Suppl. Fig 4E and F). In contrast, PINK1, a protein kinase involved in mitochondrial fission (21), and/or mitophagy (22) was higher in myotubes of both NLSDM patients (0.79 and 0.62 AU, for S-Hz and C-Hz) than in healthy controls (0.47 (0.33-0.60)), (Suppl. Fig 4G). To determine whether mitophagy contributes to altered mitochondrial oxidative capacity in myotubes of NLSDM patients, we analyzed the ratio of LC3bII/LC3bI expression in primary myotubes. LC3bII/LC3bI ratio was higher in myotubes of the S-Hz NLSDM patient (0.19 and 0.12, for S-Hz and C-Hz) compared to myotubes of healthy control subjects (0.11 (0.07-0.15)) (Suppl. Fig 4H-I) indicating increased autophagic flux and mitophagy upon ATGL deficiency due to the S-Hz mutation. Together, these differences in protein abundance may underlie the odd appearance of the mitochondria as observed by microscopy (Fig 3A and 3B), demonstrating that myotubes with dysfunctional ATGL have impaired mitochondrial dynamics.

**Figure 4.**
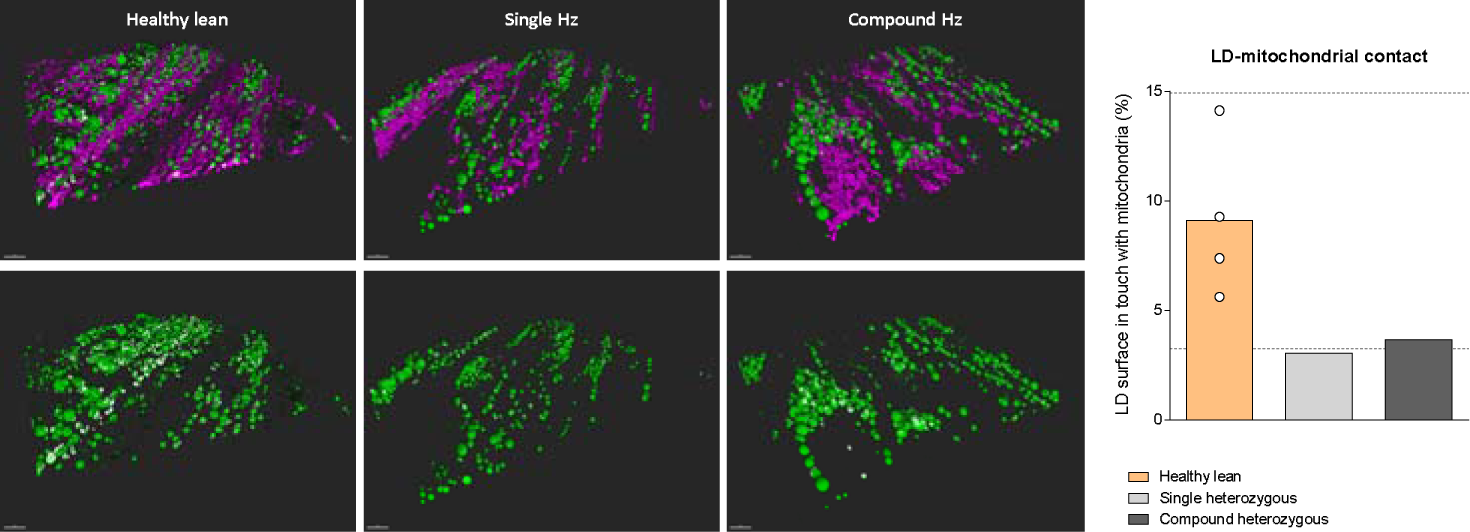
Contacts of mitochondria and lipid droplets is compromised in human primary myotubes with dysfunctional ATGL. 3-dimensional images and quantification of LD-mitochondrial contact sites in myotubes of healthy lean donors and NLSDM patients. The mitochondria were stained with TOMM20 (in magenta) and LDs were stained with Bodipy 493/503 (in green). The bottom row is showing the same LDs with the mitochondrial contact sites in white. Data are presented as mean, the 95% confidence interval of the healthy lean donors is depicted with the dashed line.

### Contacts of mitochondria and lipid droplets is compromised in human primary myotubes with dysfunctional ATGL

Contact between LDs and mitochondria was quantified as the percentage of the LD surface in contact with mitochondria. This permitted the comparison of mitochondria-LD contacts in myotubes cultured from ATGL deficient patients and healthy lean controls. The percentage of LD surface in contact with mitochondria was lower in patients with NLSDM (9.10 (3.26-14.94) vs. 3.03 vs. 3.67 % for resp. healthy lean, S-Hz and C-Hz, Fig 4).

### PPAR transcriptional activity is intact in human primary myotubes of NLSDM patients

Previously it has been shown that the lethal mitochondrial defect in the hearts of ATGL deficient mice results from defective PPAR signaling and can be rescued by PPARα agonist treatment (13). In C2C12 myotubes overexpressing ATGL PPARδ transcriptional activity was augmented (17), indicating that PPARδ agonism can compensate for ATGL deficiency in myotubes. To investigate whether primary myotubes of NLSDM patients exhibit altered PPAR transcriptional activity we analyzed PPAR-target gene expression by qRT-PCR. While mRNA expression of *PPARA, PPARD, ATGL, HSL, ADRP, CPT1, ANGPTL4,* and *PDK4* was similar among myotubes of all donors (Suppl. Fig 5A), genes involved in FA transport such as *CD36* and *FABP3* were reduced in myotubes of NLSDM patients compared to myotubes of healthy lean controls (*CD36*: 1.04 (-0.02-2.12), 0.04, and 0.07 AU for healthy lean, S-Hz and C-Hz, respectively; *FABP3:* 1.17 (0.40-1.95), 0.20, and 0.28 AU for healthy lean, S-Hz and C-Hz, respectively) (Suppl. Fig 5A). To identify whether agonists for PPARα, PPARδ or dual PPARα/δ agonist are beneficial in restoring PPAR target gene expression, myotubes of all donors were treated with either GW7647 (PPARα agonist), GW501516 (PPARδ agonist) or GFT505 (dual PPARα/δ agonist). All PPAR target genes were increasingly expressed in response to PPAR agonist treatment and to a similar extend in myotubes of NLSDM patients as in the myotubes from healthy lean donors with the exception of *PDK4*, which was more efficiently increased in myotubes of the C-Hz NLSDM patient (∼100-fold) compared to healthy lean controls and S-Hz (∼40-fold) (Suppl. Fig 5B-5D). *ATGL* gene expression was barely affected by PPARα or PPARδ activation. If any, only a modest decrease in myotubes of healthy lean donors and a modest increase in myotubes of the NLSDM patients (Suppl. Fig 5B-5D). *HSL* gene expression, on the other hand, was increased in myotubes of healthy lean controls (∼2-fold), but not in myotubes of NLSDM patients (Suppl. Fig 5B-5D). These results indicate that PPAR agonism can activate PPAR transcriptional activity even in myotubes with dysfunctional ATGL. The increase, however, does not fully restore compromised PPAR target gene expression in human primary myotubes with dysfunctional ATGL.

### PPARδ agonism increases mitochondrial respiration in myotubes with active ATGL

Next, we determined the effect of chronic PPAR-agonist treatment on mitochondrial respiration. Myotubes were incubated with a variety of PPAR-agonists during differentiation with mitochondrial function being analyzed by respirometry. In myotubes of healthy lean donors GW501516 treatment increased basal, ATP-linked, and maximal respiration by 59%, 63% and 57%, respectively (basal: 1.96 (1.18-2.73) and 3.12 (2.73-3.51) pmol/min/μg protein p<0.05, ATP-linked: 1.67 (1.02-2.32) and 2.73 (2.17-3.30) pmol/min/μg protein, p<0.01, and maximal: 3.50 (2.85-4.15) and 5.49 (5.22-5.77) pmol/min/μg protein p<0.05, for DMSO (Control) and GW501516 treated myotubes, respectively, Fig 5A-C). No increase in respiration was observed in myotubes treated with GW7647 nor GFT505 (Suppl. Fig 6A and 6B). Importantly, the positive effects of PPARδ-agonism on respiration were largely reduced in myotubes of S-Hz (+31%, +6%, and +30% in basal, ATP-linked, and maximal respiration, respectively) and absent in myotubes of C-Hz NLSDM patients (Fig 5D-F). In line with the results in the primary myotubes of NLSDM patients, ATGL inhibition reduced the effect of GW501516 on basal, ATP-linked, and maximal respiration to +14%, +24%, and +21% compared to untreated myotubes of healthy lean subjects (Fig 5G-I). Together, this suggests that active ATGL is needed for the beneficial effects of PPARδ agonism on mitochondrial respiration.

**Figure 5.**
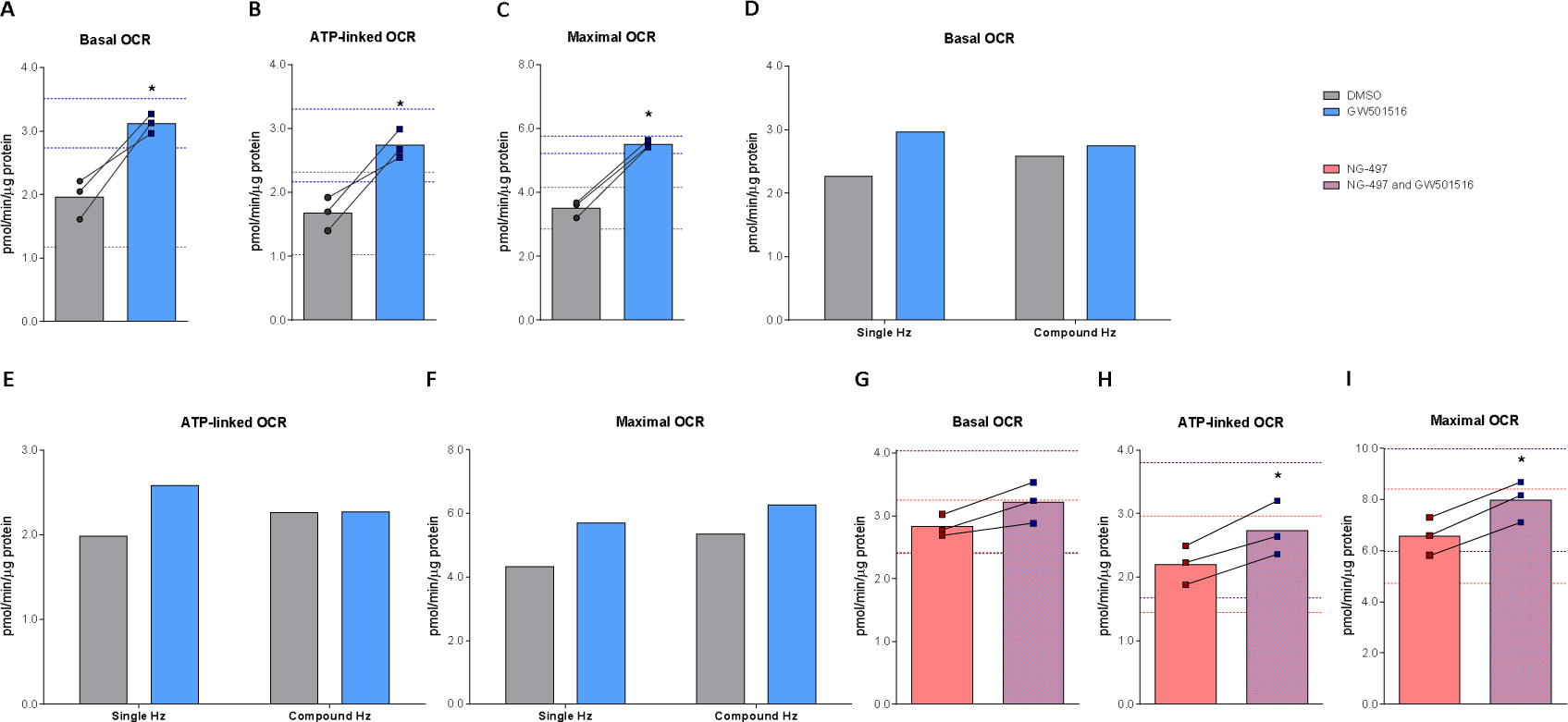
PPARδ agonism increases mitochondrial respiration in myotubes with active ATGL. (A) Basal, (B) ATP-linked and (C) maximal oxygen consumption rates of myotubes from healthy lean donors treated with GW501516. (D) Basal, (E) ATP-linked and (F) maximal oxygen consumption rates of myotubes from NLSDM patients treated with GW501516. (G) Basal, (H) ATP-linked and (I) maximal oxygen consumption rates of myotubes from healthy lean donors upon ATGL inhibition with NG-497 and treatment with GW501516. Data are presented as mean, the 95% confidence interval of the healthy lean donors is depicted with the dashed line.

**Figure 6.**
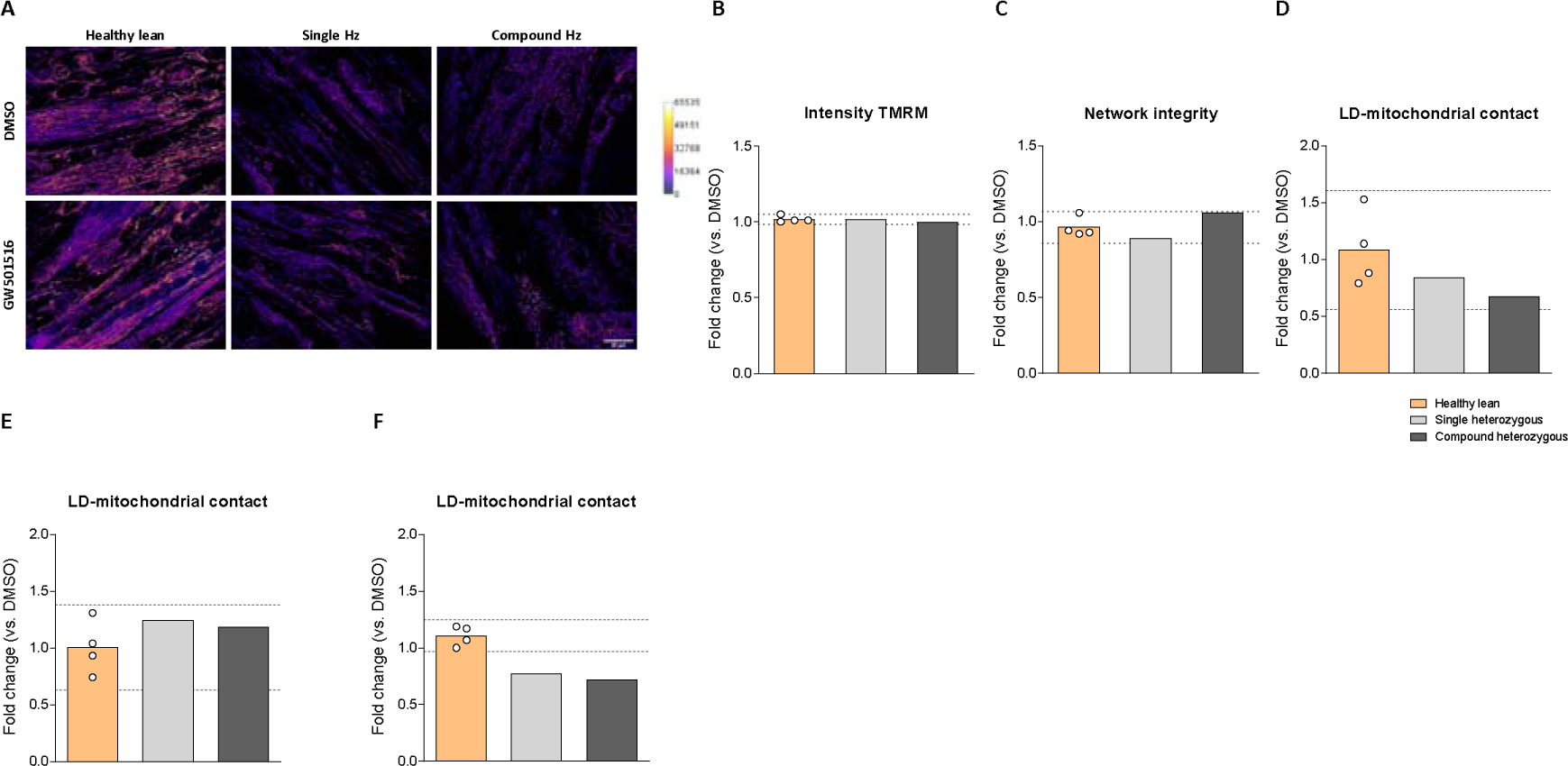
PPAR agonist treatment has no effect on mitochondrial network integrity, TMRM intensity and LD-mitochondria contact sites. (A) Spinning disk confocal images of myotubes from healthy lean donors and NLSDM patients stained for TMRM treated with GW501516. (B) Fold change of intensity TMRM and (C) network integrity upon GW501516 treatment in myotubes of healthy lean donors and NLSDM patients. Data are presented as mean, the 95% confidence interval of the healthy lean donors is depicted with the dashed line. Fold change of LD-mitochondrial contact sites in myotubes of healthy lean donors and NLSDM patients upon treatment with (D) GW501516, (E) GW7647 and (F) GFT505. Data are presented as mean, the 95% confidence interval of the healthy lean donors is depicted with the dashed line.

### PPAR agonist treatment has no effect on mitochondrial network integrity, TMRM intensity and LD-mitochondria contact sites

One of the determinants of mitochondrial oxidative capacity is mitochondrial network integrity. Next to network integrity, also the tethering of LD to mitochondria (or vice versa) affects mitochondrial respiration. Both factors contribute to mitochondrial membrane potential maintenance. In contrast to the effects of PPARδ agonist treatment on mitochondrial respiration, GW501516 had no effect on intensity of TMRM (fold change: 1.02 (0.94-1.10) vs. 1.01 vs. 1.08 for resp. healthy lean, S-Hz and C-Hz, Fig 6A and 6B). Furthermore, mitochondrial network integrity was not affected by GW501516 (fold change: 0.96 (0.86-1.07) vs. 0.89 vs. 1.06 for resp. healthy lean, S-Hz and C-Hz, Fig 6A and 6C). Similar results were found for GW7647 (Suppl. Fig 7A) and GFT505 treatment (Suppl. Fig 7B). This suggest that mitochondrial network fragmentation in ATGL deficient myotubes is independent of PPARδ activity. Furthermore, GW501516 did not affect LD-mitochondrial contact sites in primary myotubes of healthy lean donors and of NLSDM patients (fold change: 1.09 (0.56-1.61) vs. 0.84 vs. 0.67 or resp. healthy lean, S-Hz and C-Hz, Fig 6D). GW7647 treatment (fold change: 1.00 (0.63-1.38), Fig 6E) and GFT505 treatment (fold change: 1.11 (0.97-1.25), Fig 6F) did not promote the LD mitochondria contact in the myotubes of the healthy lean donors. In the NLSDM patients GW7647 did not affect LD-mitochondrial contact sites (fold change: 1.24 and 1.18 for S-Hz and C-Hz, respectively, Fig 6E). Upon GFT505 treatment the contact between LDs and mitochondria was unexpectedly reduced in both patients (fold change: 0.77 and 0.72 for S-Hz and C-Hz, respectively, Fig 6F).

## DISCUSSION

Here we used human primary myotubes from two NLSDM patients and treated human primary myotubes from healthy lean donors with an ATGL inhibitor to test the hypothesis that ATGL deficiency impairs skeletal muscle mitochondrial respiration, network connectivity and mitochondria LD contacts. Moreover, we examined if the use of PPARα and/or PPARδ agonists abolishes the functional implications of ATGL deficiency for mitochondrial function. We recapitulate *in vivo* findings of intramyocellular lipid accumulation and reduced lipolysis, and provide new data indicating impaired mitochondrial dynamics and respiration in primary myotubes from two NLSDM patients compared to healthy lean control subjects and by employing the specific human ATGL inhibitor NG-497. Mitochondrial respiration, mitochondrial network integrity and intensity TMRM were all lower in myotubes with dysfunctional ATGL or upon acute and specific inhibition of ATGL in vitro. These data suggest that for maintenance of mitochondrial network integrity, mitochondrial respiration and LD-mitochondria contacts, functional myocellular LD lipolysis via ATGL is essential. Irrespective of the phenotype of the donor, PPARδ agonist treatment improved mitochondrial respiratory capacity. This effect, however, was profoundly abolished in myotubes with dysfunctional ATGL.

We observed reduced lipolysis and increased lipid accumulation in myotubes of NLSDM patients and in response to selective ATGL inhibition (via NG-497 treatment) compared to myotubes of healthy untreated controls. ATGL-mediated TG hydrolysis provides FAs as endogenous ligands for PPARα (13, 23) and PPARδ (17) to regulate mitochondrial function. In line with this, we observed a reduced mitochondrial function in primary myotubes of NLSDM patients and upon ATGL inhibition in myotubes from healthy donors. Similarly, muscle biopsies taken from NLSDM patients exhibited reduced *ex vivo* mitochondrial function and lipid oxidative capacity compared to controls (1). In line, overexpression (17, 23) of ATGL in murine myotubes resulted in improved mitochondrial function, while global deletion of ATGL drastically reduced mitochondrial respiration in cardiac muscle (13), in cultured murine myotubes (23) and satellite cells (24). However, skeletal muscle mitochondrial function is not altered in ATGL deficient mice (25–27) and exercise intolerance of these mice rather results from diminished exogenous FA supply (25, 28, 29). In contrast, in NLSDM patients, a combination of limited exogenous FA supply (3), reduced oxidative capacity (3) and mitochondrial function (1) may contribute to exercise intolerance. To date, the mechanisms underlying the species-specific differences of ATGL deletion on skeletal muscle mitochondrial function are not known.

In global ATGL deficient mice, PPARα agonist treatment effectively restored cardiac mitochondrial dysfunction and rescued lethal cardiomyopathy (13). In contrast, PPARα agonist treatment was ineffective in improving mitochondrial function in primary myotubes of NLSDM patients and healthy donors. In NLSDM patients the pan-PPAR agonist bezafibrate (30) restored fat oxidative capacity and mitochondrial function, but was inefficient in reversing clinical symptoms such as cardiac function and muscle weakness (1). However, in primary human myotubes the dual PPARα/δ agonist was ineffective in stimulating mitochondrial function irrespective of ATGL activity. In contrast, specific PPARδ agonist treatment strongly improved mitochondrial function in myotubes from healthy donors. In line with these findings, ATGL overexpression resulted in an enhanced PPARδ, but not PPARα activity in murine myotubes (17), indicating that PPARδ may be a better target for NLSDM patients to restore mitochondrial function and improve clinical symptoms. Interestingly, PPARδ agonist treatment only improved mitochondrial respiration in myotubes with active ATGL. Similar to our results, the positive effects of PPARα agonist treatment on fat oxidation and ATP levels was blunted upon ATGL knockdown in murine myotubes (23). This suggests that in skeletal muscle ATGL activity is required to promote the positive effects of PPAR agonist treatment on mitochondrial metabolism. This may indicate that ATGL-derived fatty acids are required for the full PPAR-mediated improvements in mitochondrial respiration.

Besides impaired mitochondrial respiration, we also detected reduced intensity of TMRM and network integrity in primary myotubes of NLSDM patients and upon ATGL inhibition in myotubes of healthy controls which was unaffected by stimulation of PPAR activity. Depending on the concentration used, TMRM intensity is measured either in the quenched or the non-quenched mode (31). Here, we measured in TMRM intensity in the quenched mode, meaning that the lower intensity of TMRM upon loss of ATGL functionality indicates a higher membrane potential. In line with our results, cardiac ATGL deficiency resulted in a reduced TMRM intensity measured in the quenched mode (13). PPARα agonist treatment restored the TMRM intensity in hearts of ATGL deficient animals to wild type levels (13). Moreover, the impact of reduced ATGL activity on TMRM intensity, is reversed within hours after removal of the ATGL inhibitor. This short time span supports the notion that rather the ATGL mediated supply with FAs for oxidation than the ATGL-PPAR transcriptional activity axis is involved in maintenance of mitochondrial membrane potential in muscle. We found elevated Pink1 protein levels in both patients and an increased Lc3bII/I ratio in the S-Hz patient indicating elevated mitophagy in myotubes from NLSDM patients compared to myotubes from healthy lean controls. To remove damaged mitochondria via mitophagy, the mitochondrial network continuously undergoes cycles of fusion and fission to maintain a healthy mitochondrial network (22). Interestingly, we observed increased mitochondrial network fragmentation when ATGL was dysfunctional in myotubes indicating that ATGL activity affects mitochondria fusion/fission.

Several proteins regulate ATGL activity on a posttranslational level. Interaction with CGI-58 and G0S2 activate or inhibit ATGL, respectively (5, 32). PLIN5 regulates ATGL activity by scavenging CGI-58 when lipolysis is not activated and by releasing CGI-58 to allow ATGL activation and stimulate lipolysis. This activation of ATGL depends on the PKA mediated phosphorylation of PLIN5 (32). Cardiac muscle specific overexpression of PLIN5-S155A, a PLIN5 mutant that is not phosphorylated by PKA results in lipid accumulation, but no cardiac dysfunction (33). Similar to ATGL deficient hearts, also hearts of PLIN5-S155A transgenic mice show reduced stimulated lipolysis and impaired FA oxidation. In these hearts, however, mitochondrial function is maintained and the mitochondrial network is more fused due to reduced fission activity. This differs from our finding of enhanced fission in human primary myotubes lacking ATGL, but may be due to the differences in basal ATGL activity. While PLIN5-S155A transgenic hearts have higher ATGL protein content and *in vitro* TG hydrolase activity (33), primary myotubes from NLSDM patients have reduced or no ATGL activity. Moreover, acute ATGL inhibition also causes mitochondrial network fragmentation supporting the notion that ATGL activity is important to maintain the mitochondrial network integrity and mitochondrial quality.

Besides mitochondrial fragmentation we also observed reduced contacts between mitochondria and LDs in myotubes derived from NLSDM patients compared to healthy lean controls. As optimal FA oxidative capacity depends on FA availability and distribution across the fused mitochondrial network (19), it is suggested that mitochondria interacting with LDs have a higher ATP production capacity (18). Accordingly, starvation induced ATGL-mediated TG hydrolysis and FA oxidation is associated with a tighter connection between LDs and mitochondria (19). PPAR agonist treatment had no effect on mitochondria-LD contacts, indicating that the ATGL-PPAR transcriptional activity does not play a role in the connection between these organelles. Together, a disbalanced mitochondrial network and reduced connections between LDs and mitochondria contribute to a reduced respiration in myotubes of NLSDM patients.

Besides its impact on respiration, a disbalance in mitochondrial fusion and fission is also associated with muscle atrophy (34–37) and muscle weakness (34, 35). Myopathic symptoms observed in NLSDM patients are muscle weakness (1, 4) which, in the more advanced cases, is accompanied by muscle atrophy (4). Mutations in OPA1 cause autosomal-dominant optical atrophy and late onset myopathy in patients (38) and reduced mitochondrial fusion in muscles of mutant rats (38). While Opa1 or Mfn2 protein content was similar in myotubes with dysfunctional ATGL compared to healthy lean controls, we observed a profound reduction in Mfn1 protein abundance in myotubes of NLSDM patients. Similarly, knockdown of Mfn1, but not Mfn2, resulted in less mitochondrial fusion events and chronic alcohol feeding induced myopathy in rats and was associated with reduced Mfn1, but not Mfn2 and Opa1 levels. In addition, mitochondrial fusion was impaired and mitochondrial energy metabolism was compromised (38). Recently Yue et al. 2022 (24) showed that satellite cell specific ATGL deficiency impairs muscle regeneration after muscle damage and regenerated muscles have reduced fiber size. In line, we found myotube differentiation was compromised in NLSDM patients and when ATGL was inhibited in healthy cells. These results suggests that the myopathy observed in NLSDM patients may be due to a reduced muscle regenerative capacity potentially as a consequence of a disbalance in mitochondrial fusion and fission activity and mitochondrial dysfunction.

In conclusion, dysfunctional ATGL compromises mitochondrial dynamics, reduces mitochondrial-LD contacts and decreases respiratory capacity. Since defects in mitochondrial function are linked to myopathy our data suggest that myopathy in NLSDM patients also results from a compromised respiratory capacity and a fragmented mitochondrial network. PPARδ agonist treatment increases oxygen consumption rate only in the presence of active ATGL and does not restore mitochondrial function in myotubes lacking ATGL, pointing towards alternative treatment regimens to ameliorate myopathy in patients with NLSDM.

## METHODS

### Donor details

Satellite cells were isolated from needle muscle biopsies taken from the m. vastus lateralis from a patient with NLSDM and a heterozygous mutant carrier participated in a previous study (1). The NLSDM patient had a compound heterozygous (C-Hz) mutation causing a frame-shift at amino acid position 270 and a missense mutation 584C → T causing a proline to leucine change at amino acid 195. The heterozygous mutant carrier (single heterozygous, S-Hz) was a carrier of the frame-shift at amino acid position 270. Four young healthy lean donors were used as controls (39, 40).

### Human primary myotube culture

Satellite cells were isolated from muscle biopsies and immunopurified using a CD56 antibody (5.1H11, Developmental Studies Hybridoma Bank, Iowa City, IA, USA) as described previously (41). Myoblasts were grown at 37°C in a humidified atmosphere with 5% CO2 in DMEM (1 g/L glucose, Gibco, Thermo Fisher Scientific, Waltham, MA, USA) supplemented with 15% Fetal Bovine Serum (FBS, Invitrogen-Life Technologies, Eugene, OR, USA), 0.5 mg/ml BSA (A6003, Sigma, Zwijndrecht, the Netherlands), 0.5 mg/ml fetuin (10344-026, Gibco, Thermo Fisher Scientific, Waltham, MA, USA), 10 ng/ml human epidermal growth factor (PHG0311, Invitrogen-Life Technologies, Eugene, OR, USA), 1 µM dexamethasone (D8893, Sigma, Zwijndrecht, the Netherlands), 50 µg/ml gentamycin (15750-037, Invitrogen-Life Technologies, Eugene, OR, USA), 1.25 µg/ml fungizone (15290-018, Gibco, Thermo Fisher Scientific, Waltham, MA, USA) and 25 U/ml penicillin/streptomycin (15070-063, Invitrogen-Life Technologies, Eugene, OR, USA). After reaching ∼80% confluence cells were plated in 6-wells, 12-wells, 8-wells µ-Slide (80826, Ibidi GmbH, Martinsried, Germany), 8-well removable chamber slides (80841, Ibidi GmbH, Martinsried, Germany) or 96-well plates (for Seahorse, Agilent Technologies, Santa Clara, CA, USA). At 90-100% confluency differentiation was initiated by changing the medium to α-MEM Glutamax (Gibco, Thermo Fisher Scientific, Waltham, MA, USA) supplemented with 2% FBS, 0.5 mg/ml fetuin and 100 U/ml penicillin/streptomycin and changed every other day. For cellular respiration measurements, human primary myotubes were differentiated without any antibiotics. After 4-6 days of differentiation satellite cells were fully differentiated and experiments were performed. For the experiments examining the effects of PPAR-agonist treatment and/or ATGL inhibition, PPAR-agonists and NG-497 were added to the differentiation medium for the whole differentiation period, and medium was changed every day. Human primary myotubes were treated with PPARα agonist GW7647 (1 µM; G6793, Sigma Zwijndrecht, the Netherlands), PPARδ agonist GW501516 (1 µM; SML1491, Sigma Zwijndrecht, the Netherlands) or with the dual PPAR α/δ agonist GFT505 (175 nM, B1030-5, BioVision, Waltham, MA, USA). In addition, human primary myotubes from the healthy lean donors were treated with the human ATGL inhibitor NG-497(20) (40 µM). DMSO served as vehicle control.

### Electron microscopy

Biopsy material was fixed with 2% paraformaldehyde and 0.1% glutaraldehyde at 4°C. Ultrathin cryosections were cut under liquid nitrogen and were stained with uranyl acetate for contrast. Sections were imaged using a FEI T12 transmission electron microscopy coupled to a FEI Eagle CCD camera.

### Intramyocellular lipid accumulation and myotube quality

Myoblasts were grown on coverslips in a 12-wells plate. Upon full differentiation myotubes from healthy lean donors were incubated with 50 μM oleic acid with or without 40 μM NG-497 to acutely inhibit ATGL for ∼16 hours. Subsequently, human primary myotubes were fixed as described for the lipolysis experiments. LDs were stained using Bodipy 493/503 (D3922, Molecular Probes, Leiden, The Netherlands; diluted 1:100) and nuclei with DAPI (1:100 dilution). To analyze myotube differentiation, sections were stained using antibodies against sarcomeric myosin (MF20, Developmental Studies Hybridoma Bank, Iowa City, IA, USA) and α-actinin (a653, Fürst/Potsdamm). Subsequently sections were incubated with appropriate AlexaFluor-conjugated secondary antibodies (Thermo Fisher Scientific, Waltham, MA, USA) and DAPI. Images were acquired using a Nikon E800 fluorescent microscope (Nikon, Amsterdam, the Netherlands) coupled to a Nikon DS-Fi1c color CCD camera (Nikon, Amsterdam, the Netherlands) using the appropriate filters.

### Respirometry

Respiration rates were measured using the Seahorse XF96 extracellular flux analyzer (Agilent Technologies, Santa Clara, CA, USA). Before analysis human primary myotubes were incubated 1 hour at 37°C in unbuffered XF assay medium adjusted to pH7.4 with 1 M NaOH. XF assay medium for glucose oxidation consisted of 8.308 mg/ml DMEM 5030 supplemented with 116 nM NaCl, 5.5 mM glucose, 4 mM GlutaMax and 1 mM sodium pyruvate. Respiration rates were measured in quadruplicates with a ∼7-minute interval. Basal respiration was determined followed by injection of oligomycin (1 µM final concentration) to measure leak respiration. Subsequently, the uncoupler FCCP (1 µM final concentration) was injected to measure maximal uncoupled respiration. To determine non-mitochondrial linked respiration complex I and III inhibitors rotenone and antimycin A (both 1 µM final concentration) were injected. Respiration rates data was corrected for total protein content using either the BCA protein assay (Thermo Fisher Scientific, Waltham, MA, USA) or DC protein assay (Bio-Rad, Veenendaal, the Netherlands).

### Lipolysis

Human primary myotubes were grown and differentiated in 8-well removable chamber slides. Upon full differentiation myotubes were incubated overnight (+16 hours) with 200 µM oleic acid and 2 µM Bodipy-FL labelled C16 fatty acids (D3821, Thermo Fisher Scientific, Waltham, MA, USA). After an overnight pulse-(timepoint 0h) and after a 3-hour chase period in DMEM supplemented with 0.1 mM glucose and 0.5% essential fatty acid free BSA, cells were fixated with 3.7% formaldehyde in PBS for 10 minutes After fixation nuclei were stained with DAPI diluted 1:100 in PBS and mounted with Mowiol and covered with #1.5 coverslips and stored in the dark before imaging.

### Mitochondrial networks

Human primary myotubes were grown and differentiated in 8-wells µ-Slides. Upon full differentiation myotubes were incubated for 30 minutes with 250 nM tetramethyorhadamine methyl ester perchlorate (TMRM) (T668, Thermo Fisher Scientific, Waltham, MA, USA), Hoechst 34580 (1 mg/ml; diluted 1:1000; H21486, Molecular Probes, Leiden, the Netherlands) and CellMask Deep Red diluted (1:2000; C10046, Invitrogen-Life Technologies, Eugene, OR, USA) in DMEM without phenol red supplemented with 5.5 mM glucose, 2% FBS, and 2 mM GlutaMAX (35050061, Gibco, Thermo Fisher Scientific, Waltham, MA, USA) to visualize mitochondria, nuclei and plasma membrane respectively. Subsequently myotubes were washed 3 times with pre-warmed DPBS (A12856-01, Gibco, Thermo Fisher Scientific, Waltham, MA, USA). For live-cell spinning disk imaging, DPBS was exchanged by DMEM supplemented with 5.5 mM glucose, 2% FBS, and 2 mM GlutaMAX, and 25 mM HEPES.

### Lipid droplet – mitochondria contact sites

Human primary myotubes were grown and differentiated on a glass coverslip in 12-wells plate. On the last day of differentiation, myotubes were incubated with 200 µM oleic acid overnight (+16h) to induce LD formation. After oleic acid incubation myotubes were fixated with cold 3.7% formaldehyde in PBS for 10 minutes and permeabilized with 0.1% TX-100 in PBS for 5 minutes. Mitochondria were visualized using an antibody against TOMM20 (ab56783, Abcam, Cambridge, UK; diluted 1:50 in 0.05% Tween/PBS; 45 minutes at room temperature). Thereafter, myotubes were washed and incubated with AlexaFluor568 Goat anti Rabbit secondary antibody (A11011; Invitrogen, Groningen, The Netherlands; 1:200 in 0.05% Tween/PBS), Bodipy 493/503 (1:100), and DAPI (1:100). After washing, the coverslips were removed from the 12-wells plate and mounted with Mowiol on an object glass slide and stored in the dark before imaging.

### Confocal laser scanning microscopy and image analysis

#### Lipolysis

Images were acquired on a Leica TCS SP8 confocal microscope using a 63× oil immersion/1.4 N.A objective using 2048×2048 pixels resulting in a 90 nm x 90 nm pixel size. For deconvolution purposes z-stacks of 5 slices with a step size of 0.15 µm were acquired. DAPI and Bodipy-FL were excited using respectively the 405 nm and 488 nm laser lines. Emission was collected with PMT detectors between 415-460 nm and 500-540 nm for respectively DAPI and Bodipy-FL. The pinhole was set at 1 Airy unit. Images were deconvolved with Huygens Professional Software (Scientific Volume Imaging B.V., Hilversum, the Netherlands) and analyzed with ImageJ (42). From the Bodipy-FL images binary images were created to quantify LD content. LD content was corrected for cell number (based on DAPI staining; multinucleated cells were taken into account). Basal lipolysis was determined by subtracting LD content after 3 hours from timepoint 0 measurement.

#### Lipid droplet – mitochondria contact sites

Complete z-stacks were acquired with a Leica TCS SP8 confocal microscope using a 63× oil immersion/1.4 N.A. objective with a 0.10 µm z-step. A 3× optical zoom was used and images consisted of 1024×1024 pixels resulting in a 60 nm x 60 nm pixel size. DAPI, Bodipy 493/503 and TOMM20-AlexaFluor568 were excited using 405 nm, 488 nm and 594 nm laser lines, respectively, and emission was collected between 415-460 nm, 500-540 nm and 590-655 nm, respectively. Emission for DAPI was collected with a hybrid detector and gating, i.e. detecting photons with a certain lifetime, was set at 0.3-6.0 ns. For Bodipy 493/503 and TOMM20-AlexaFluor586 emission was collected using a PMT detector. A pinhole of 1 Airy unit was used. Images were deconvolved using Huygens Professional Software and analyzed with ImageJ (42). To determine LD-mitochondria contact sites z-stacks were analyzed for percentage of LD surface in touch with mitochondria and number of contact sites per LD. The LD channel was first filtered using a Gaussian filter. A binary image was created and a watershed algorithm was used to segment LDs. Subsequently boundaries were labeled using the MorphoLibJ plugin (43). The channel with mitochondria was filtered using a white top hat filter and a binary image was created. LD-mitochondria contact sites were determined by creating a new image which depicted only the pixels that were present in the LD boundaries image and the thresholded image of the mitochondria. The Analyze Regions 3D command from the MorphoLibJ plugin was used to analyze the volume of LD boundaries and contact sites. Based on these volumes the percentage of the LD surface in touch with mitochondria was calculated. In addition, LD number and total contact sites were determined to calculate the average number of contact sites per LD.

### Live-cell spinning disk confocal microscopy and image analysis

Human primary myotubes were imaged with a FEI CorrSight spinning disk confocal microscope, equipped with an ORCA-Flash4.0 V2 CMOS camera. Imaging was performed at 37°C and 5% CO_2_ with a 63× oil immersion 1.4 N.A. objective (Zeiss). The camera was cropped at 1656×1228 pixels resulting in a pixel size of 103 x 103 nm. Z-stacks of 75-90 slices were acquired with a step size of 0.17 µm at 15 random positions per well. The CellMask Deep Red staining was used to manually set the starting point of each z-stack using the 640 nm laser. TMRM and Hoechst 34580 were excited with respectively 405 nm and 561 nm laser lines, and emission was collected through the Semrock quadband bandpass filter (FF01-446-523-600-677-25). Images were deconvolved using Huygens Professional Software. Matlab R2021a (The MathWorks, Inc., Natick, MA, USA) was used to analyze images for total mitochondrial volume, number of mitochondrial network particles and TMRM intensity. Camera noise was subtracted from images and images were filtered using a top-hat spatial filter and median filter. A binary image was created to form a mask of the image with mitochondria. For each image mitochondrial network particle size, total mitochondrial volume, and TMRM intensity was determined. From these data the mitochondrial fragmentation index (MFI) was determined by dividing the number of mitochondrial network particles by total mitochondrial volume. A higher MFI indicates a more fragmented mitochondrial network.

### Western blotting

Human primary myotubes were grown and differentiated in a 12-well plate. Myotubes were collected either using 150 µl Bioplex lysis buffer (171304011, Bio-Rad, Veenendaal, the Netherlands) supplemented with 2 mM PMSF, or RIPA-buffer (Supplemental Table 1). Samples were subjected to SDS-polyacrylamide gel electrophoresis. Different pre-cast polyacrylamide gels were used depending on the protein of interest (Suppl table 1). For the 4-12 Bolt^TM^ Bis-Tris gels (NW04125, Thermo Fisher Scientific, Waltham, MA, USA) equal protein loading was determined by staining the gels with Instant blue (ab119211, LI-COR Biosciences, Westburg, Leusden, The Netherlands) after gel electrophoresis.

Stained gels were scanned using the Odyssey CLx infrared detector, and Image Studio^TM^ Software (LI-COR Biosciences, Westburg, Leusden, the Netherlands) was used to quantify optical density for calculating equal protein loading. Proteins were transferred onto a nitrocellulose membrane using the Trans-Blot Turbo system (Bio-Rad, Veenendaal, the Netherlands) and stained with Revert^TM^ 700 (926-11011, LI-COR Biosciences, Westburg, Leusden, The Netherlands) to check for efficient protein transfer. When using TGX Stain-Free gels, equal volumes of protein samples were loaded and gels were scanned using the Gel Doc^TM^ EZ Imager (Bio-Rad, Veenendaal, the Netherlands) to correct for differences in protein loading. After protein transfer, nitrocellulose membranes were scanned using the Gel Doc^TM^ EZ Imager to check for efficient protein transfer. To detect specific proteins, primary antibodies against ATGL, CGI-58, HSL, OXPHOS antibody cocktail (detecting all 5 mitochondrial electron transport chain complexes), PGC1-α, Mfn1, Mfn2, OPA1, DRP1, Fis1, PINK1, and LC3b were used. A list of antibodies is provided in supplemental Table 1. Appropriate HRP-conjugated secondary antibodies were used for ECL detection using the ChemicDoc (Bio-Rad, Veenendaal, the Netherlands), and appropriate IRDye680-conjugated or IRDye800-conjugated secondary antibodies were used for detection using the Odyssey CLx infrared detector.

### Real-time (RT) quantitative (q)-PCR

For RNA isolation from human primary myotubes cells were harvested in Trizol® Reagent (15596026, Thermo Fisher Scientific, Waltham, MA, USA) and stored at -80°C. Total RNA was isolated using the RNeasy Mini spin columns (74104, Qiagen, Germantown, MD, USA). cDNA was synthesized from the isolated RNA samples using the High-Capacity RNA-to-cDNA^TM^ kit (4387406, Applied Biosystems^TM^, Waltham, MA, USA) according to the manufacturer protocol. Subsequently RT-qPCR reactions were performed using the CFX384 Touch Real-Time PCR Detection System (Bio-Rad, Veenendaal, the Netherlands) and the following parameters: 10 min 50°C, 10 min 95°C followed by 40 cycles of 15 sec 95°C and 1 min 60°C. Gene expression data were normalized to the geometric mean of the housekeeping genes *RPLP0* and *RPL26*. Primers, probes and TaqMan^TM^ assays used are shown in Supplemental Table 2.

### RT-qPCR for mtDNA copy number

Human primary myotubes were washed with ice-cold DPBS. Subsequently, cells were scraped in DPBS and collected in 15 mL conical tubes. After centrifugation for 5 minutes at room temperature, the pellet was resuspended in 200 μL ice-cold DPBS and stored at -80°C. DNA was isolated using the DNeasy kit (69504, Qiagen, Germantown, MD, USA) according to the manufacturers protocol. RT-qPCR was performed using the following parameters: 2 min 50°C, 10 min 95°C followed by 40 cycles of 15 sec 95°C and 1 min 60°C. Primers and probes used are shown in Supplemental Table 2. Ct-values were used to calculate DNA concentrations.

### Statistics

Results for the healthy lean donors are presented as mean (95% CI). Since the NLSDM patients have different ATGL mutations and these mutations have different consequence for disease severity *in vivo,* data from these human primary myotubes are presented individually. Data from the NLSDM patients were considered as different from the healthy lean when these datapoints were below or above the 95% confidence interval of the mean of the healthy lean donors. Data on ATGL inhibition or PPAR agonist treatment effects were either shown as fold change when lacking any effects or as control vs. treatment. A paired t-test was performed in SPSS version 28 (IBM Corp., Armonk, NY, USA) to test for significance and p<0.05 was considered as statistically significant.

### Study approval

The studies were approved by the institutional medical committee of Maastricht University and participants signed an informed consent prior to participation (1, 39, 40).

### Data availability

Data underlying the findings described in this manuscript may be available upon request by the corresponding author.

## AUTHOR CONTRIBUTIONS

AG, RZ, MS and MKCH designed the study. AG, TvdW, GS, EK, KK and MS conducted experiments and acquired data. AG, MS and MKCH interpret data and wrote manuscript. GFG provided reagents. TvdW, GS, GFG, EK, KK and RZ revised the manuscript critically and approved final version of the manuscript.

## ACKNOWLEDGEMENTS

The work of AG and MKCH is partly supported by the Netherlands Cardiovascular Research Initiative: an initiative with support of the Dutch Heart Foundation (CVON2014-02 ENERGISE).

---

**Supplemental Table 1.**
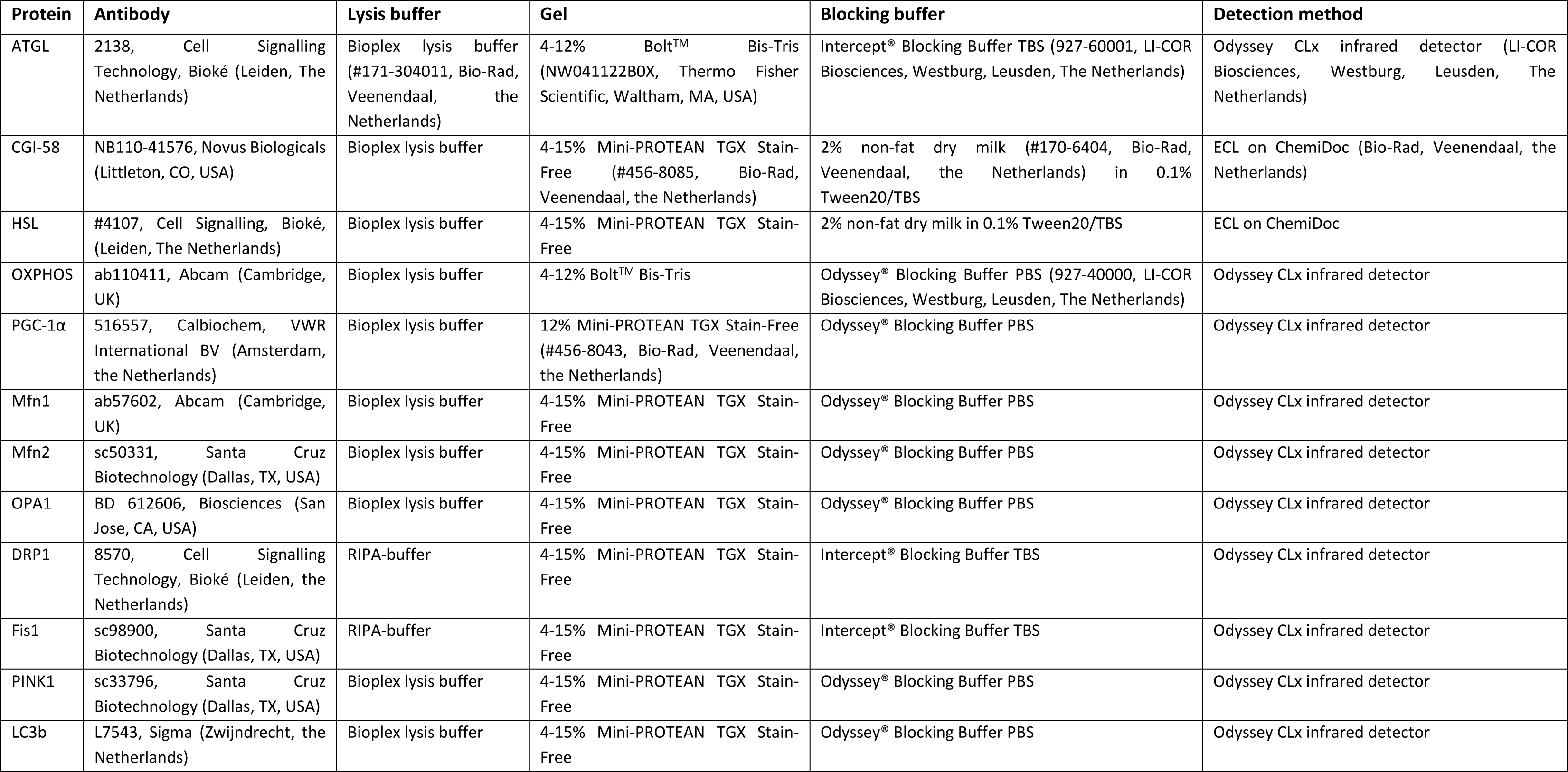
Methods Western Blotting.

**Supplemental Table 2.**
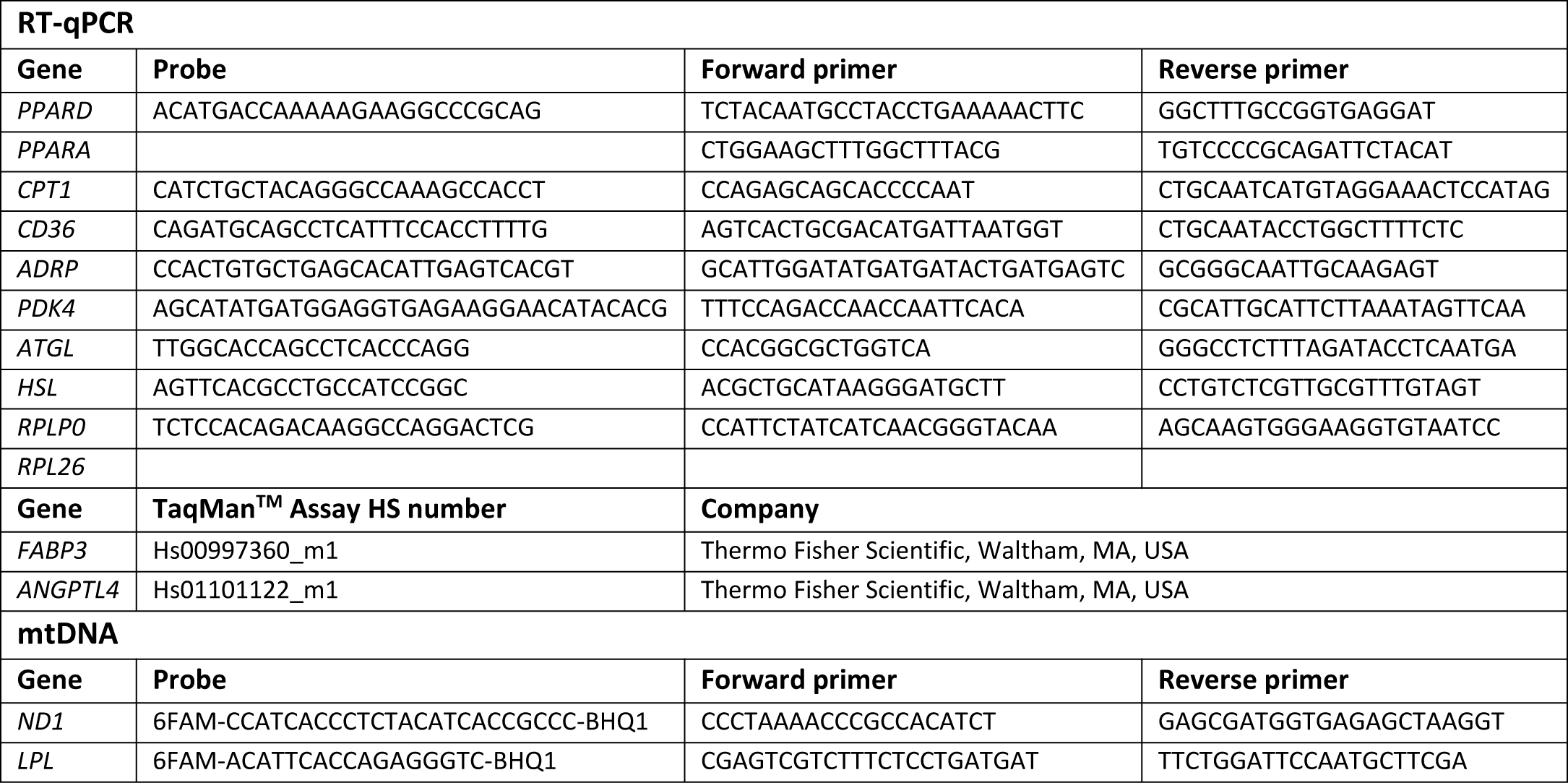
Probes, primer and TaqMan^TM^ assays RT-qPCR.

## SUPPLEMENTAL FIGURE LEGENDS

**Supplemental Figure 1. Intramyocellular lipid content and mitochondrial abnormalities in biopsies of muscle with dysfunctional ATGL.** Transmission electron microscopy images of skeletal muscle biopsies of NLSDM patients. Besides high intramyocellular lipid content, mitochondrial arrangement is abnormal. In the endurance trained athlete mitochondria are located on both sides of the z-line, while this is not the case for the mitochondria of the NLSDM patients. In addition, mitochondria are also dispersed longitudinally in large groups between the contractile elements.

**Supplemental Figure 2. ATGL deficiency compromises myotube differentiation.** (A) Myotubes of healthy lean donors and NLSDM patients stained for MF20 (red), α-actinin (green) and DAPI (blue). (B) Myotubes of healthy lean donors upon ATGL inhibition with NG-497.

**Supplemental Figure 3. The effect of ATGL inhibition on intensity TMRM and network integrity is acute.** (A) Spinning disk confocal images of TMRM staining myotubes from healthy lean donors in which ATGL was inhibited with NG-497 during differentiation, but imaged when the ATGL inhibitor was removed for ∼3.5 hours. (B) Spinning disk confocal images of TMRM staining myotubes from healthy lean donors upon acute ATGL inhibition with NG-497.

**Supplemental Figure 4. Protein content of the different players in mitochondrial dynamics.** Western blot analysis of (A) PGC-1α, (B) MFN1, (C) MFN2, (D) OPA1, (E) DRP1, (F) FIS1, (G) PINK1, (H) LC3b-I (cytosolic) and LC3b-II (autophagosome)), and (I) ratio LC3b-II/LC3b-I. Data are presented as mean, the 95% confidence interval of the healthy lean donors is depicted with the dashed line.

**Supplemental Figure 5. Gene expression.** (A) Gene expression of PPARs, PPAR target genes and genes involved in lipolysis in myotubes of healthy lean donors and NLSDM patients, and fold change upon treatment with (B) GW501516, (C) GW7647 and (D) GFT505.

**Supplemental Figure 6. Changes in mitochondrial respiratory capacity upon PPAR agonist treatment.** Fold change in oxygen consumption rates upon (A) GW7647 and (B) GFT505 treatment in myotubes of healthy lean donors and NLSDM patients.

**Supplemental Figure 7. Intensity TMRM and network integrity upon PPAR agonist treatment.** Spinning disk confocal images of TMRM stained mitochondria, and fold change of intensity TMRM and network integrity of myotubes from healthy lean donors and NLSDM patients upon treatment with (A) GW7647 and (B) GFT505. Data are presented as mean, the 95% confidence interval of the healthy lean donors is depicted with the dashed line.

## REFERENCES

1. van de Weijer T, Havekes B, Bilet L, Hoeks J, Sparks L, Bosma M, et al. Effects of bezafibrate treatment in a patient and a carrier with mutations in the PNPLA2 gene, causing neutral lipid storage disease with myopathy. Circ Res. 2013;112(5):e51–4.

2. Fischer J, Lefevre C, Morava E, Mussini JM, Laforet P, Negre-Salvayre A, et al. The gene encoding adipose triglyceride lipase (PNPLA2) is mutated in neutral lipid storage disease with myopathy. Nat Genet. 2007;39(1):28–30.

3. Laforet P, Orngreen M, Preisler N, Andersen G, and Vissing J. Blocked muscle fat oxidation during exercise in neutral lipid storage disease. Arch Neurol. 2012;69(4):530–3.

4. Pennisi EM, Arca M, Bertini E, Bruno C, Cassandrini D, D’Amico A, et al. Neutral Lipid Storage Diseases: clinical/genetic features and natural history in a large cohort of Italian patients. Orphanet J Rare Dis. 2017;12(1):90.

5. Schreiber R, Xie H, and Schweiger M. Of mice and men: The physiological role of adipose triglyceride lipase (ATGL). Biochim Biophys Acta Mol Cell Biol Lipids. 2019;1864(6):880–99.

6. Missaglia S, Maggi L, Mora M, Gibertini S, Blasevich F, Agostoni P, et al. Late onset of neutral lipid storage disease due to novel PNPLA2 mutations causing total loss of lipase activity in a patient with myopathy and slight cardiac involvement. Neuromuscul Disord. 2017;27(5):481–6.

7. Tavian D, Maggi L, Mora M, Morandi L, Bragato C, and Missaglia S. A novel PNPLA2 mutation causing total loss of RNA and protein expression in two NLSDM siblings with early onset but slowly progressive severe myopathy. Genes Dis. 2021;8(1):73–8.

8. Goodpaster BH, He J, Watkins S, and Kelley DE. Skeletal muscle lipid content and insulin resistance: evidence for a paradox in endurance-trained athletes. J Clin Endocrinol Metab. 2001;86(12):5755–61.

9. Krssak M, Falk Petersen K, Dresner A, DiPietro L, Vogel SM, Rothman DL, et al. Intramyocellular lipid concentrations are correlated with insulin sensitivity in humans: a 1H NMR spectroscopy study. Diabetologia. 1999;42(1):113–6.

10. Forouhi NG, Jenkinson G, Thomas EL, Mullick S, Mierisova S, Bhonsle U, et al. Relation of triglyceride stores in skeletal muscle cells to central obesity and insulin sensitivity in European and South Asian men. Diabetologia. 1999;42(8):932–5.

11. Bergman BC, Perreault L, Strauss A, Bacon S, Kerege A, Harrison K, et al. Intramuscular triglyceride synthesis: importance in muscle lipid partitioning in humans. Am J Physiol Endocrinol Metab. 2018;314(2):E152–E64.

12. de la Rosa Rodriguez MA, and Kersten S. Regulation of lipid droplet-associated proteins by peroxisome proliferator-activated receptors. Biochim Biophys Acta Mol Cell Biol Lipids. 2017;1862(10 Pt B):1212–20.

13. Haemmerle G, Moustafa T, Woelkart G, Buttner S, Schmidt A, van de Weijer T, et al. ATGL-mediated fat catabolism regulates cardiac mitochondrial function via PPAR-alpha and PGC-1. Nat Med. 2011;17(9):1076–85.

14. Tanaka T, Yamamoto J, Iwasaki S, Asaba H, Hamura H, Ikeda Y, et al. Activation of peroxisome proliferator-activated receptor delta induces fatty acid beta-oxidation in skeletal muscle and attenuates metabolic syndrome. Proc Natl Acad Sci U S A. 2003;100(26):15924–9.

15. Feng YZ, Nikolic N, Bakke SS, Boekschoten MV, Kersten S, Kase ET, et al. PPARdelta activation in human myotubes increases mitochondrial fatty acid oxidative capacity and reduces glucose utilization by a switch in substrate preference. Arch Physiol Biochem. 2014;120(1):12–21.

16. Zizola C, Kennel PJ, Akashi H, Ji R, Castillero E, George I, et al. Activation of PPARdelta signaling improves skeletal muscle oxidative metabolism and endurance function in an animal model of ischemic left ventricular dysfunction. Am J Physiol Heart Circ Physiol. 2015;308(9):H1078–85.

17. Meex RC, Hoy AJ, Mason RM, Martin SD, McGee SL, Bruce CR, et al. ATGL-mediated triglyceride turnover and the regulation of mitochondrial capacity in skeletal muscle. Am J Physiol Endocrinol Metab. 2015;308(11):E960–70.

18. Bleck CKE, Kim Y, Willingham TB, and Glancy B. Subcellular connectomic analyses of energy networks in striated muscle. Nat Commun. 2018;9(1):5111.

19. Rambold AS, Cohen S, and Lippincott-Schwartz J. Fatty acid trafficking in starved cells: regulation by lipid droplet lipolysis, autophagy, and mitochondrial fusion dynamics. Dev Cell. 2015;32(6):678–92.

20. Grabner GF, Guttenberger N, Mayer N, Migglautsch-Sulzer AK, Lembacher-Fadum C, Fawzy N, et al. Small-Molecule Inhibitors Targeting Lipolysis in Human Adipocytes. J Am Chem Soc. 2022;144(14):6237–50.

21. Han H, Tan J, Wang R, Wan H, He Y, Yan X, et al. PINK1 phosphorylates Drp1(S616) to regulate mitophagy-independent mitochondrial dynamics. EMBO Rep. 2020;21(8):e48686.

22. Dahlmans D, Houzelle A, Schrauwen P, and Hoeks J. Mitochondrial dynamics, quality control and miRNA regulation in skeletal muscle: implications for obesity and related metabolic disease. Clin Sci (Lond). 2016;130(11):843–52.

23. Biswas D, Ghosh M, Kumar S, and Chakrabarti P. PPARalpha-ATGL pathway improves muscle mitochondrial metabolism: implication in aging. FASEB J. 2016;30(11):3822–34.

24. Yue F, Oprescu SN, Qiu J, Gu L, Zhang L, Chen J, et al. Lipid droplet dynamics regulate adult muscle stem cell fate. Cell Rep. 2022;38(3):110267.

25. Dube JJ, Sitnick MT, Schoiswohl G, Wills RC, Basantani MK, Cai L, et al. Adipose triglyceride lipase deletion from adipocytes, but not skeletal myocytes, impairs acute exercise performance in mice. Am J Physiol Endocrinol Metab. 2015;308(10):E879–90.

26. Sitnick MT, Basantani MK, Cai L, Schoiswohl G, Yazbeck CF, Distefano G, et al. Skeletal muscle triacylglycerol hydrolysis does not influence metabolic complications of obesity. Diabetes. 2013;62(10):3350–61.

27. Nunes PM, van de Weijer T, Veltien A, Arnts H, Hesselink MK, Glatz JF, et al. Increased intramyocellular lipids but unaltered in vivo mitochondrial oxidative phosphorylation in skeletal muscle of adipose triglyceride lipase-deficient mice. Am J Physiol Endocrinol Metab. 2012;303(1):E71–81.

28. Huijsman E, van de Par C, Economou C, van der Poel C, Lynch GS, Schoiswohl G, et al. Adipose triacylglycerol lipase deletion alters whole body energy metabolism and impairs exercise performance in mice. Am J Physiol Endocrinol Metab. 2009;297(2):E505–13.

29. Schoiswohl G, Schweiger M, Schreiber R, Gorkiewicz G, Preiss-Landl K, Taschler U, et al. Adipose triglyceride lipase plays a key role in the supply of the working muscle with fatty acids. J Lipid Res. 2010;51(3):490–9.

30. Willson TM, Brown PJ, Sternbach DD, and Henke BR. The PPARs: from orphan receptors to drug discovery. J Med Chem. 2000;43(4):527–50.

31. Perry SW, Norman JP, Barbieri J, Brown EB, and Gelbard HA. Mitochondrial membrane potential probes and the proton gradient: a practical usage guide. Biotechniques. 2011;50(2):98–115.

32. Sanders MA, Madoux F, Mladenovic L, Zhang H, Ye X, Angrish M, et al. Endogenous and Synthetic ABHD5 Ligands Regulate ABHD5-Perilipin Interactions and Lipolysis in Fat and Muscle. Cell Metab. 2015;22(5):851–60.

33. Kolleritsch S, Kien B, Schoiswohl G, Diwoky C, Schreiber R, Heier C, et al. Low cardiac lipolysis reduces mitochondrial fission and prevents lipotoxic heart dysfunction in Perilipin 5 mutant mice. Cardiovasc Res. 2020;116(2):339–52.

34. Favaro G, Romanello V, Varanita T, Andrea Desbats M, Morbidoni V, Tezze C, et al. DRP1-mediated mitochondrial shape controls calcium homeostasis and muscle mass. Nat Commun. 2019;10(1):2576.

35. Tezze C, Romanello V, Desbats MA, Fadini GP, Albiero M, Favaro G, et al. Age-Associated Loss of OPA1 in Muscle Impacts Muscle Mass, Metabolic Homeostasis, Systemic Inflammation, and Epithelial Senescence. Cell Metab. 2017;25(6):1374–89 e6.

36. Romanello V, and Sandri M. The connection between the dynamic remodeling of the mitochondrial network and the regulation of muscle mass. Cell Mol Life Sci. 2021;78(4):1305–28.

37. Dulac M, Leduc-Gaudet JP, Reynaud O, Ayoub MB, Guerin A, Finkelchtein M, et al. Drp1 knockdown induces severe muscle atrophy and remodelling, mitochondrial dysfunction, autophagy impairment and denervation. J Physiol. 2020;598(17):3691–710.

38. Eisner V, Lenaers G, and Hajnoczky G. Mitochondrial fusion is frequent in skeletal muscle and supports excitation-contraction coupling. J Cell Biol. 2014;205(2):179–95.

39. Vosselman MJ, Hoeks J, Brans B, Pallubinsky H, Nascimento EB, van der Lans AA, et al. Low brown adipose tissue activity in endurance-trained compared with lean sedentary men. Int J Obes (Lond). 2015;39(12):1696–702.

40. Grevendonk L, Connell NJ, McCrum C, Fealy CE, Bilet L, Bruls YMH, et al. Impact of aging and exercise on skeletal muscle mitochondrial capacity, energy metabolism, and physical function. Nat Commun. 2021;12(1):4773.

41. Sparks LM, Moro C, Ukropcova B, Bajpeyi S, Civitarese AE, Hulver MW, et al. Remodeling lipid metabolism and improving insulin responsiveness in human primary myotubes. PLoS One. 2011;6(7):e21068.

42. Schneider CA, Rasband WS, and Eliceiri KW. NIH Image to ImageJ: 25 years of image analysis. Nature methods. 2012;9(7):671–5.

43. Legland D, Arganda-Carreras I, and Andrey P. MorphoLibJ: integrated library and plugins for mathematical morphology with ImageJ. Bioinformatics. 2016;32(22):3532–4.

