## Supplemental materials for "ATGL-mediated lipolysis is essential for myocellular mitochondrial function and augments PPARδ-induced improvements in mitochondrial respiration"

**Supplemental Table 1. Methods Western Blotting.**

| Protein | Antibody | Lysis buffer | Gel | Blocking buffer | Detection method |
|---|---|---|---|---|---|
| ATGL | 2138, Cell Signalling Technology, Bioké (Leiden, The Netherlands) | Bioplex lysis buffer (#171-304011, Bio-Rad, Veenendaal, the Netherlands) | 4-12% Bolt™ Bis-Tris (NW041122B0X, Thermo Fisher Scientific, Waltham, MA, USA) | Intercept® Blocking Buffer TBS (927-60001, LI-COR Biosciences, Westburg, Leusden, The Netherlands) | Odyssey CLx infrared detector (LI-COR Biosciences, Westburg, Leusden, The Netherlands) |
| CGI-58 | NB110-41576, Novus Biologicals (Littleton, CO, USA) | Bioplex lysis buffer | 4-15% Mini-PROTEAN TGX Stain-Free (#456-8085, Bio-Rad, Veenendaal, the Netherlands) | 2% non-fat dry milk (#170-6404, Bio-Rad, Veenendaal, the Netherlands) in 0.1% Tween20/TBS | ECL on ChemiDoc (Bio-Rad, Veenendaal, the Netherlands) |
| HSL | #4107, Cell Signalling, Bioké, (Leiden, The Netherlands) | Bioplex lysis buffer | 4-15% Mini-PROTEAN TGX Stain-Free | 2% non-fat dry milk in 0.1% Tween20/TBS | ECL on ChemiDoc |
| OXPHOS | ab110411, Abcam (Cambridge, UK) | Bioplex lysis buffer | 4-12% Bolt™ Bis-Tris | Odyssey® Blocking Buffer PBS (927-40000, LI-COR Biosciences, Westburg, Leusden, The Netherlands) | Odyssey CLx infrared detector |
| PGC-1α | 516557, Calbiochem, VWR International BV (Amsterdam, the Netherlands) | Bioplex lysis buffer | 12% Mini-PROTEAN TGX Stain-Free (#456-8043, Bio-Rad, Veenendaal, the Netherlands) | Odyssey® Blocking Buffer PBS | Odyssey CLx infrared detector |
| Mfn1 | ab57602, Abcam (Cambridge, UK) | Bioplex lysis buffer | 4-15% Mini-PROTEAN TGX Stain-Free | Odyssey® Blocking Buffer PBS | Odyssey CLx infrared detector |
| Mfn2 | sc50331, Santa Cruz Biotechnology (Dallas, TX, USA) | Bioplex lysis buffer | 4-15% Mini-PROTEAN TGX Stain-Free | Odyssey® Blocking Buffer PBS | Odyssey CLx infrared detector |
| OPA1 | BD 612606, Biosciences (San Jose, CA, USA) | Bioplex lysis buffer | 4-15% Mini-PROTEAN TGX Stain-Free | Odyssey® Blocking Buffer PBS | Odyssey CLx infrared detector |
| DRP1 | 8570, Cell Signalling Technology, Bioké (Leiden, the Netherlands) | RIPA-buffer | 4-15% Mini-PROTEAN TGX Stain-Free | Intercept® Blocking Buffer TBS | Odyssey CLx infrared detector |
| Fis1 | sc98900, Santa Cruz Biotechnology (Dallas, TX, USA) | RIPA-buffer | 4-15% Mini-PROTEAN TGX Stain-Free | Intercept® Blocking Buffer TBS | Odyssey CLx infrared detector |
| PINK1 | sc33796, Santa Cruz Biotechnology (Dallas, TX, USA) | Bioplex lysis buffer | 4-15% Mini-PROTEAN TGX Stain-Free | Odyssey® Blocking Buffer PBS | Odyssey CLx infrared detector |
| LC3b | L7543, Sigma (Zwijndrecht, the Netherlands) | Bioplex lysis buffer | 4-15% Mini-PROTEAN TGX Stain-Free | Odyssey® Blocking Buffer PBS | Odyssey CLx infrared detector |

**Supplemental Table 2. Probes, primer and TaqMan$^{TM}$ assays RT-qPCR.**

| **RT-qPCR** | | | |
|---|---|---|---|
| **Gene** | **Probe** | **Forward primer** | **Reverse primer** |
| *PPARD* | ACATGACCAAAAAGAAGGCCCGCAG | TCTACAATGCCTACCTGAAAAACTTC | GGCTTTGCCGGTGAGGAT |
| *PPARA* | | CTGGAAGCTTTGGCTTTACG | TGTCCCCGCAGATTCTACAT |
| *CPT1* | CATCTGCTACAGGGCCAAAGCCACCT | CCAGAGCAGCACCCCAAT | CTGCAATCATGTAGGAAACTCCATAG |
| *CD36* | CAGATGCAGCCTCATTTCCACCTTTTG | AGTCACTGCGACATGATTAATGGT | CTGCAATACCTGGCTTTTCTC |
| *ADRP* | CCACTGTGCTGAGCACATTGAGTCACGT | GCATTGGATATGATGATACTGATGAGTC | GCGGGCAATTGCAAGAGT |
| *PDK4* | AGCATATGATGGAGGTGAGAAGGAACATACACG | TTTCCAGACCAACCAATTCACA | CGCATTGCATTCTTAAATAGTTCAA |
| *ATGL* | TTGGCACCAGCCTCACCCAGG | CCACGGCGCTGGTCA | GGGCCTCTTTAGATACCTCAATGA |
| *HSL* | AGTTCACGCCTGCCATCCGGC | ACGCTGCATAAGGGATGCTT | CCTGTCTCGTTGCGTTTGTAGT |
| *RPLP0* | TCTCCACAGACAAGGCCAGGACTCG | CCATTCTATCATCAACGGGTACAA | AGCAAGTGGGAAGGTGTAATCC |
| *RPL26* | | | |
| **Gene** | **TaqMan$^{TM}$ Assay HS number** | **Company** | |
| *FABP3* | Hs00997360_m1 | Thermo Fisher Scientific, Waltham, MA, USA | |
| *ANGPTL4* | Hs01101122_m1 | Thermo Fisher Scientific, Waltham, MA, USA | |
| **mtDNA** | | | |
| **Gene** | **Probe** | **Forward primer** | **Reverse primer** |
| *ND1* | 6FAM-CCATCACCCTCTACATCACCGCCC-BHQ1 | CCCTAAAACCCGCCACATCT | GAGCGATGGTGAGAGCTAAGGT |
| *LPL* | 6FAM-ACATTCACCAGAGGGTC-BHQ1 | CGAGTCGTCTTTCTCCTGATGAT | TTCTGGATTCCAATGCTTCGA |

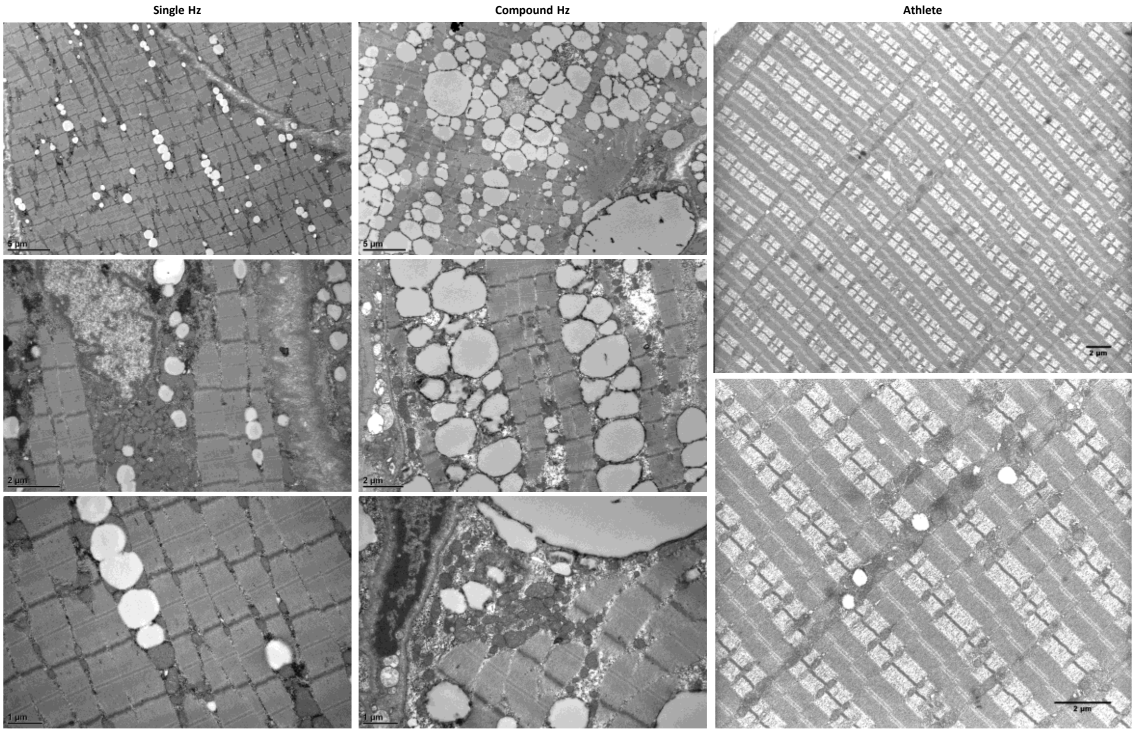


**Supplemental Figure 1. Intramyocellular lipid content and mitochondrial abnormalities in biopsies of muscle with dysfunctional ATGL.** Transmission electron microscopy images of skeletal muscle biopsies of NLSDM patients. Besides high intramyocellular lipid content, mitochondrial arrangement is abnormal. In the endurance trained athlete mitochondria are located on both sides of the z-line, while this is not the case for the mitochondria of the NLSDM patients. In addition, mitochondria are also dispersed longitudinally in large groups between the contractile elements.

**A**

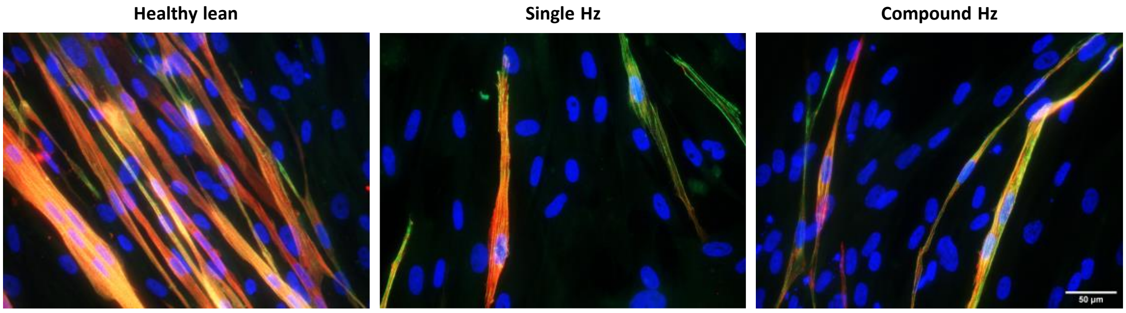


**B**

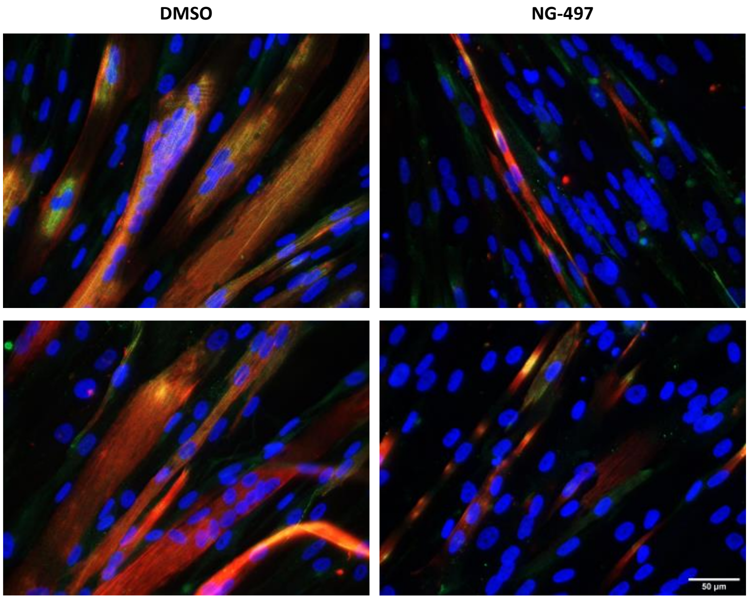


**Supplemental Figure 2. ATGL deficiency compromises myotube differentiation.** (A) Myotubes of healthy lean donors and NLSDM patients stained for MF20 (red), α-actinin (green) and DAPI (blue). (B) Myotubes of healthy lean donors upon ATGL inhibition with NG-497.

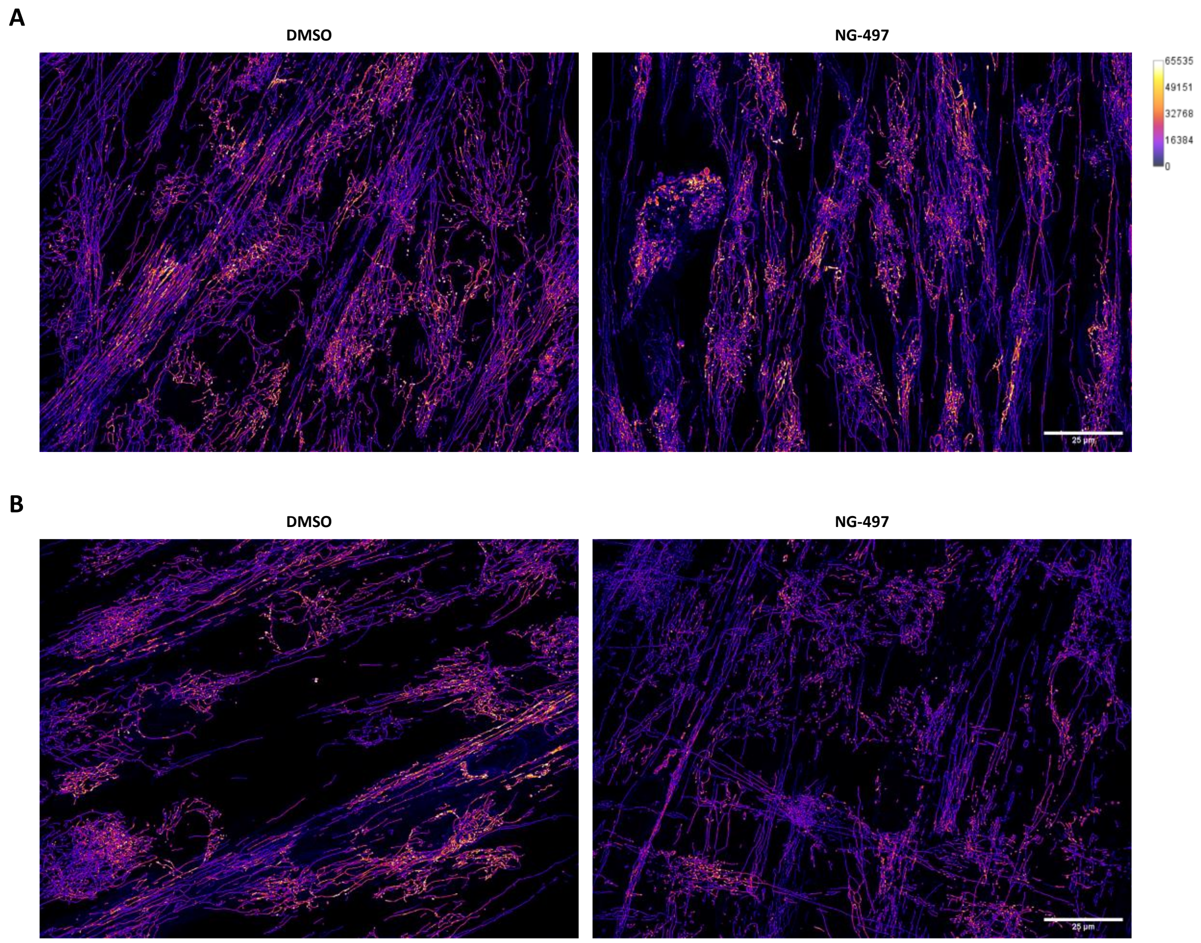


**Supplemental Figure 3. The effect of ATGL inhibition on intensity TMRM and network integrity is acute.** (A) Spinning disk confocal images of TMRM staining myotubes from healthy lean donors in which ATGL was inhibited with NG-497 during differentiation, but imaged when the ATGL inhibitor was removed for ~3.5 hours. (B) Spinning disk confocal images of TMRM staining myotubes from healthy lean donors upon acute ATGL inhibition with NG-497.

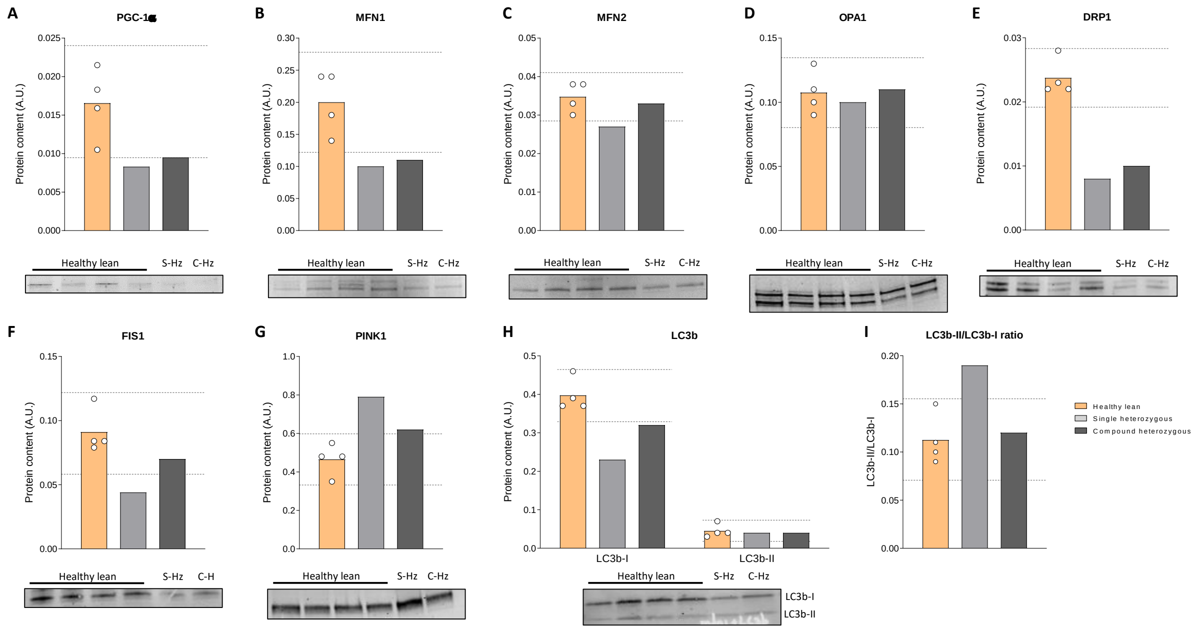


**Supplemental Figure 4. Protein content of the different players in mitochondrial dynamics.** Western blot analysis of (A) PGC-1α, (B) MFN1, (C) MFN2, (D) OPA1, (E) DRP1, (F) FIS1, (G) PINK1, (H) LC3b-I (cytosolic) and LC3b-II (autophagosome) ), and (I) ratio LC3b-II/LC3b-I. Data are presented as mean, the 95% confidence interval of the healthy lean donors is depicted with the dashed line.

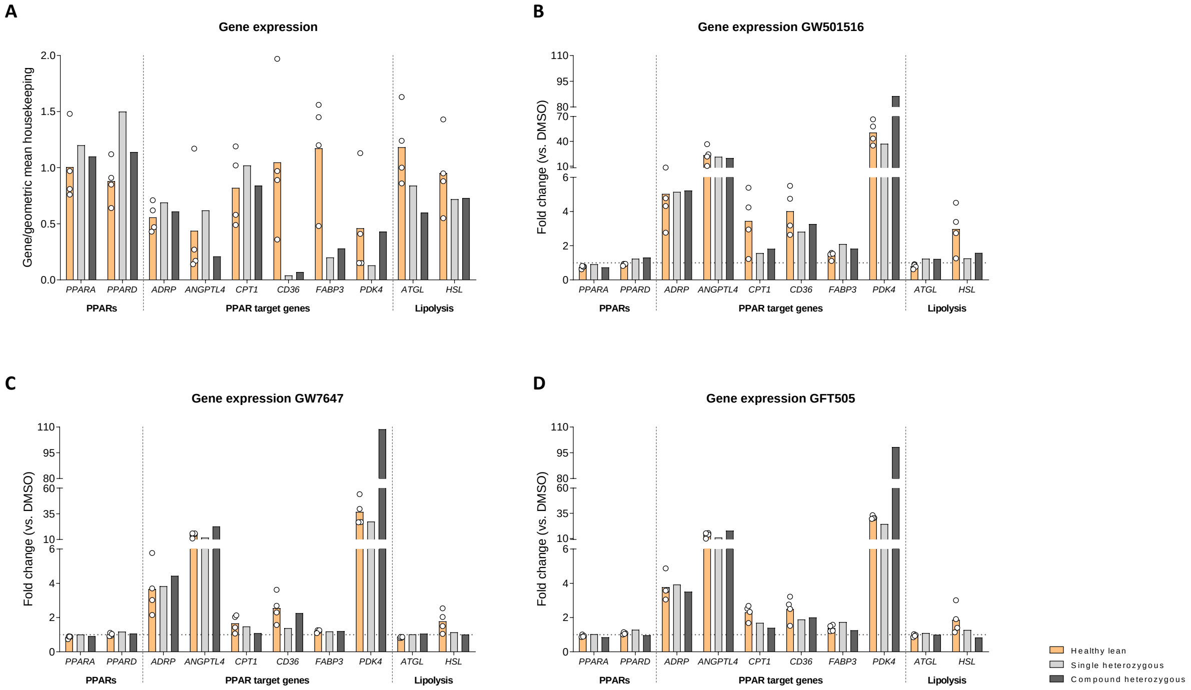


**Supplemental Figure 5. Gene expression.** (A) Gene expression of PPARs, PPAR target genes and genes involved in lipolysis in myotubes of healthy lean donors and NLSDM patients, and fold change upon treatment with (B) GW501516, (C) GW7647 and (D) GFT505.

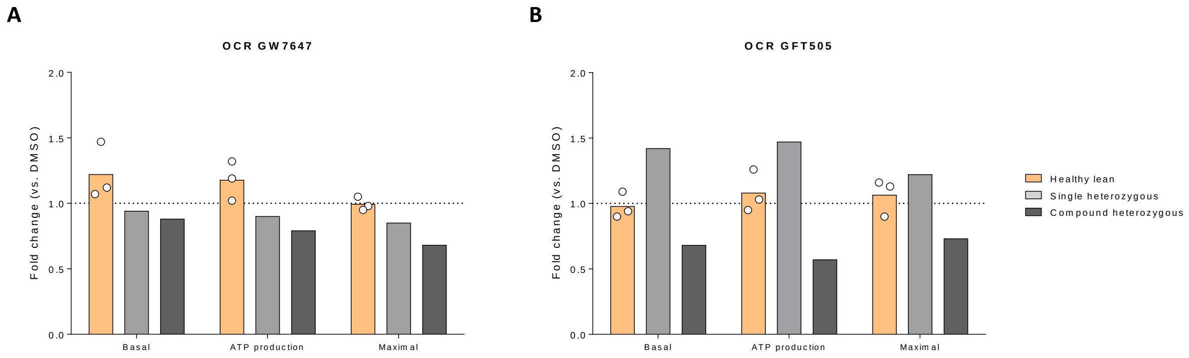


**Supplemental Figure 6. Changes in mitochondrial respiratory capacity upon PPAR agonist treatment.** Fold change in oxygen consumption rates upon (A) GW7647 and (B) GFT505 treatment in myotubes of healthy lean donors and NLSDM patients.

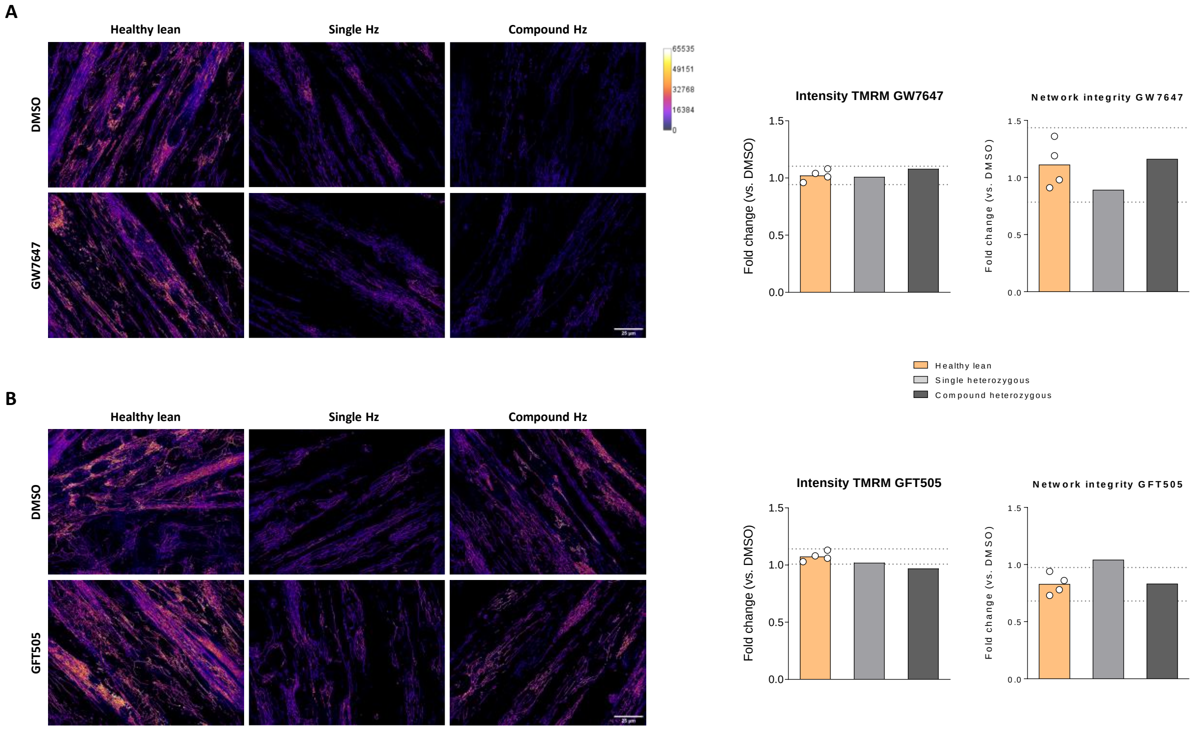


**Supplemental Figure 7. Intensity TMRM and network integrity upon PPAR agonist treatment.** Spinning disk confocal images of TMRM stained mitochondria, and fold change of intensity TMRM and network integrity of myotubes from healthy lean donors and NLSDM patients upon treatment with (A) GW7647 and (B) GFT505. Data are presented as mean, the 95% confidence interval of the healthy lean donors is depicted with the dashed line.
